# Comprehensive drug efficacy data for mucinous ovarian carcinoma using a novel and extensive biobank of patient-derived organoid models

**DOI:** 10.64898/2026.04.06.716848

**Authors:** Olivia Craig, Carolina Salazar, Suad Abdirahman, Nikita Dalvi, Niveditha Rajadevan, Jennii Luu, Robert Vary, Susanne Ramm, Karla J Cowley, Ratana Lim, Briony Milesi, Christopher Tagkalidis, Patricia Wojtowicz, Stuart Bencraig, Genevieve Dall, Maree Pechlivanis, Breana G Galea, My Fitzgerald, Rita Saoud, Minh A To, Richard Lupat, Song Li, Catherine Kennedy, Rachel Delahunty, Prue Allan, Robert G Ramsay, Alicia Oshlack, Anna DeFazio, Clare L Scott, Kaylene J Simpson, Orla M McNally, Kylie L Gorringe

**Affiliations:** Peter MacCallum Cancer Centre, 305 Grattan St, Melbourne, VIC 3052, Australia; The University of Melbourne, Parkville, VIC 3052, Australia; The Royal Women’s Hospital, Parkville, VIC 3052, Australia; The Walter and Eliza Hall Institute of Medical Research, Parkville, VIC 3052, Australia; Centre for Cancer Research, The Westmead Institute for Medical Research, Sydney, NSW, 2145, Australia; Department of Gynaecological Oncology, Westmead Hospital, Sydney, NSW 2145, Australia; Faculty of Medicine and Health, The University of Sydney, Sydney, NSW 2006, Australia; The Daffodil Centre, The University of Sydney, and Cancer Council NSW, NSW 2006, Australia; The Mercy Hospital for Women and Austin Health, Heidelberg VIC 3084, Australia; Barwon Health and Deakin University, Geelong VIC 3220, Australia

**Author notes:** Equal contribution (Olivia Craig, Carolina Salazar). These authors contributed equally.

## Abstract

Mucinous Ovarian Carcinoma (MOC) is a rare ovarian cancer histological subtype with distinct pathology, genomics and clinical outcomes compared to other epithelial ovarian cancers. Accordingly, there is little evidence to guide clinical care, particularly in the use of systemic therapies, and the field has lacked informative and diverse pre-clinical models. We developed MOC-specific methods for generating tumour organoids with a success rate of 70% for long-term cultured lines (n=19). Organoid lines were developed from localised, advanced and recurrent tumours, including from biopsy tissue, and represent diverse genomic features not previously captured by existing cell lines. The organoid lines were highly similar to the tumours of origin for genomic and immunohistochemical markers. Screening using a panel of 11 chemotherapy agents highlighted resistance to standard-of-care agents such as carboplatin. Gastrointestinal cancer chemotherapy agents and their combination regimens lacked activity. Paclitaxel was often highly potent at low doses but failed to kill all cells. However, less frequently used drugs such as gemcitabine, topotecan and doxorubicin inhibited many of the lines more effectively than paclitaxel. Available, but non-standard of care, chemotherapy agents should be considered for clinical management of MOC. This is the largest (by ∼10 fold) cohort of fully characterised patient-derived MOC organoid lines described and the first with extensive drug screening data affording an opportunity for drug discovery and screening for personalised treatment.

## Introduction

Mucinous Ovarian Carcinoma (MOC) is a rare histotype of epithelial ovarian cancer that has a poor prognosis when diagnosed at a late stage, with a median survival of only 12 – 33 months for patients with advanced stage disease [1]. It is distinct from the more common High Grade Serous Ovarian Carcinoma (HGSOC), and only more recent molecular and histological classification has allowed it to be accurately distinguished from mucinous extra-ovarian metastases masquerading as primary ovarian tumours [2]. Due to disease rarity and diagnostic difficulties, human clinical trial data for MOC is extremely limited and challenging to collect. For the same reasons, few verified pre-clinical models have been established from these tumours, with scarce patient derived xenograft (PDX) models developed [3,4]. Available cell lines are similarly limited, with MCAS [5], RMUG-S [6] and JHOM-1 (RIKEN Cell Bank RCB:1676) the most representative of the key molecular events and likely truly of MOC origin [7]. Such a limited inventory fails to capture the heterogeneity of this rare histotype, with features such as *ERBB2* amplification, found in up to 27% of MOC tumours [8], not represented. As such, research in this field has moved slowly. Patients with MOC have poor tumour responses to standard of care ovarian cancer platinum-based chemotherapy regimens like carboplatin/paclitaxel [9-11], but little evidence has been generated to support the use of other therapeutics. Consequently, MOC patient survival has not improved over time [12].

For decades, cell lines have been the cornerstone of *in vitro* biomedical research. However, in recent years tumour organoids have emerged as a potentially superior pre-clinical model system, reported to more accurately reflect parent tumours [13,14]. Organoids are formed from human or animal tissue samples that have been enriched for stem-like cell populations [15,16]. These populations are embedded in a synthetic gel designed to closely mimic the native basement membrane and cultured with a cocktail of growth factors specific to the physiological niche of the originating tissue. The combination of these elements enables the stem-like cell populations to proliferate and self-organise into 3D structures that approximate the complexity of the architecture, and in turn function, of the corresponding organ or tissue [17].

Since the first reported cultures [18], organoids have been used to model a range of cancer types, including colon, lung, breast, brain and oesophageal cancers [19]. More recently, ovarian cancer organoids have been generated across the range of epithelial histotypes, including a few mucinous tumour organoid lines, demonstrating their faithful phenotypic and molecular reflection of parent tumours [20,21]. There have been sparse reports of one or two MOC organoid lines within larger ovarian cancer organoid generation projects [22,23], but these tend to be overshadowed by the focus on the more common HGSOC, with limited functional data generated using these models.

Here, we describe the first focused effort to develop an extensive and representative biobank of MOC organoids for use in testing therapies. As MOC is thought to develop along a continuum of progression through benign and borderline neoplasms [8,24], we also collected benign and borderline mucinous ovarian tumours for organoid generation on the basis of forming relevant precursor models.

## Methods

### Organoid generation from human tissue samples

Human tumour samples were obtained from patients at Australian hospitals undergoing surgical cytoreduction or biopsy (Supplementary Table 1). All patients provided informed consent for the use of their tissue for research. The study was approved by the Human Research Ethics Committee at Peter MacCallum Cancer Centre (HREC #20/34).

Fresh tissues were collected into cold DMEM/F-12 containing HEPES, GlutaMAX, penicillin and streptomycin (Organoid wash solution, Supplementary Methods) and transported to the lab for timely processing. Tissue collected at interstate sites was cryopreserved in 10% dimethylsulphoxide (DMSO)/90% fetal calf serum (FCS) and shipped to the central lab on dry ice. Size permitting, samples were divided into sections, with the majority reserved for organoid derivation and one piece snap frozen in liquid nitrogen, one piece processed into a formalin-fixed paraffin-embedded (FFPE) block and one piece cryopreserved in Recovery™ Cell Culture Freezing Medium (Gibco).

The tissue designated for organoid generation was transferred to one well of a six-well plate and minced using sterile surgical scissors before the addition of ∼1 mL digestion media (2.5 mg/mL Collagenase Type II with 10 μM Y-27632 in DMEM/F-12). The butt of a 1 mL syringe was used to mash the pieces into the bottom of the well, before transfer to a 15 mL tube using a cut transfer pipette and up to another 14 mL digestion media to wash the well. The tube containing tissue pieces was placed on a shaking platform and gently rocked while incubated at 37 °C for 15-30 min. The contents of the tube were further broken up using a cut transfer pipette before being passed through a 70 μm cell strainer. Filtered solution was centrifuged at 500 x *g* at 4 °C for 10 min and if resultant pellet was visibly red, a red blood cell digestion was performed for 5 min using Red Blood Cell Lysis Buffer (Roche) and stopped with addition of DMEM/F-12. Following centrifugation, the remaining pellet was resuspended in 0.1-1 mL Matrigel (Corning Phenol Red Free Lot#923-100) and 25 μL drops were plated into wells of a 48-well plate (Greiner). Plates were incubated for 20 min at 37 °C before the addition of our optimised medium MOC-ORGM3 (Supplementary Methods) or Kopper et al [25] organoid growth medium.

### Organoid culture

Organoids were cultured with either MOC-ORGM3 or Kopper medium (Supplementary Methods) that was completely refreshed every 2–3 days. When organoids reached high density, they were scraped from wells using pipette tips and cold DMEM/F-12 to wash and combine wells. Organoids were centrifuged at 500 x g at 4 °C for 10 min and pellet resuspended in TrypLE Express (Gibco). Organoids were incubated at 37 °C for 10–20 min with manual trituration performed as necessary, before TrypLE Express was diluted with cold DMEM/F-12 and cells were centrifuged again. The resultant pellet was resuspended in Matrigel (Corning) with at least a 1:3 dilution and 25 μL drops were plated into a 48-well plate (Greiner). Plates were incubated for 20 min at 37 °C before the addition of either MOC-ORGM3 or Kopper organoid growth media.

### Organoid cryopreservation and recovery

To determine optimal recovery after cryopreservation, we used a robust organoid line (ORG38) frozen at two different timepoints: during exponential growth phase and when at high density. We compared various conditions including dissociation into cell clusters using TrypLE Express and maintaining intact organoids within the Matrigel domes. Five cryopreservatives were tested, including ThermoFisher Cell Recovery Solution, CryoStor CS10 (Stem Cell Technologies), organoid growth medium supplemented with 10% DMSO, FCS supplemented with 10% DMSO, and 1:1 growth media and FCS supplemented with 10% DMSO. To freeze whole organoids, Matrigel containing organoids was lifted off the plate and transferred into a cryovial in relevant cryopreservative. To generate cell clusters, organoids from two confluent domes were processed as described above and transferred to a cryovial containing the appropriate cryopreservative. After freezing overnight at -80 °C in a Mr Frosty container (ThermoFisher Scientific), vials were transferred to liquid nitrogen for three months. Organoids were recovered as described below and images taken weekly.

Following the optimisation experiment, organoids were cryopreserved as follows: 2–3 days before organoids were predicted to be at high density, media was removed from wells to be cryopreserved and replaced with Cell Recovery Solution. Organoids were gently rotated on ice for 30–60 min before wells were pooled with cold DMEM/F-12 and centrifuged at 500 x *g* at 4° C for 10 min. Supernatant was discarded and organoids were resuspended in Cell Recovery freezing media (for early passage organoids) or ORGM3/10% DMSO and transferred to cryovials before freezing at -80 °C prior to transfer to liquid nitrogen.

For recovery, cryopreserved vials were retrieved from liquid nitrogen and quickly thawed in a water bath. Contents of cryovials were immediately diluted dropwise with warm organoid media then transferred into 15 mL conical tubes and further diluted with up to 7 mL pre-warmed organoid wash solution or media, before centrifuging at 500 x *g* at 4 °C for 10 min. The resultant pellet was resuspended in an appropriate volume of Matrigel depending on the size of the pellet and 25 μL drops were plated into each well of a 48-well plate (Greiner). Plates were incubated for 20 min at 37 °C before the addition of either ORGM3 or Kopper media.

### Organoid growth factor dependency testing

Organoids were dissociated to single cells (described above) and 20 μL domes containing 1500–3000 cells were seeded in 48-well plates. ORGM3 media was added, either complete or lacking specific elements including: 50% Forskolin, no Wnt3A, no Noggin, no RSPO, no FGF10, and no Wnt3A/Noggin/RSPO with 3-5 replicate wells for each condition. Post seeding, brightfield imaging was performed weekly at 4X on the Cytation C10 Cell imaging multi-mode reader (BioTek, Agilent) with a maximum projection of a stack of three z-heights as described previously [26]. Passaging was performed on all wells when the organoids became confluent in the complete media wells. Organoids were counted using automated detection of objects using QuPath on the brightfield images.

### Immunohistochemistry (IHC) characterisation

Matrigel domes containing large organoids prior to passaging were scraped from wells using pipette tips with cold DMEM/F-12 and combined into a pre-wet 1.5 mL LoBind® tube (Eppendorf). Organoids were centrifuged at 500 x *g* at 4 °C for 10 min and gently but quickly resuspended in 100 μL 1% liquid agarose before transfer to a cryomold. Organoid-agarose mixture was left to cool and set before fixing in 10% Neutral Buffered Formalin for 24 h and processing into a paraffin block.

Matching organoid and parent-tumour FFPE blocks were sectioned at 4 μm before IHC of key MOC markers CK7, CK20, PAS, PAX8, p53 and HER2 was performed using clinically validated antibodies by the Anatomical Pathology Department, Peter MacCallum Cancer Centre (Supplementary Table 2). CK7, CK20 and PAX8 were scored as: a) positive: >90% cells all strongly staining; b) patchy positive: >10% of cells staining with variable intensity; c) focal positive <10% of cells staining; d) negative: no cells staining. HER2 was scored as per the gastric cancer guidelines: 0: <10%; 1+: Incomplete weak staining in >10%; 2+: weak complete staining or moderate incomplete staining in >10%; 3+: strong complete staining in >10%. TP53 staining was scored abnormal positive, wild-type and abnormal negative as previously described [27]. For concordance analysis, exactly matching categories were “concordant”. Categories that could potentially be explained by tumour heterogeneity, for example focal positive and patchy positive, or 1+ and 2+, were called “partial concordance”. “Discordance” was determined when one sample was positive and another negative.

### DNA/RNA extraction

#### Organoids

DNA: 4–5 wells of confluent organoids were combined into a single pre-wet 1.5 mL LoBind® tube using cold DMEM/F-12 and pelleted by centrifugation at 500 x *g* at 4 °C for 10 min. The supernatant was discarded and DNA lysis buffer (100 mM Tris-CI pH 8, 50 mM EDTA pH 8, 100 mM NaCI and 1% SDS) and 20 μL Proteinase K (Qiagen) were added. Samples were incubated at 56 °C for 4 h, with brief vortexing every hour. 6 M NaCI was added to each sample and mixed before incubating on ice for 10 min and centrifuging at 20,000 x g for 10 min. The supernatant was transferred to a fresh LoBind® tube and the DNA precipitated with 20 mg/mL Glycogen (Roche) and 100% ethanol. Tubes were incubated at room temperature for 10 min before being centrifuged at 20,000 x *g* for 10 min. 70% ethanol was mixed with the pellets by vortexing. Samples were centrifuged at 20,000 x *g* for 5 min and the supernatant removed before being air dried for 10 min to evaporate remaining ethanol. Pellets were resuspended in 30–50 μL TE buffer and vortexed prior to overnight incubation at 4 °C. DNA was quantified using the Qubit fluorometer and reagents (ThermoFisher Scientific).

RNA: 4–5 wells of confluent organoids were combined into a single pre-wet 1.5 mL LoBind® tube using cold DMEM/F-12 and pelleted by centrifugation at 500 x *g* at 4 °C for 10 min. Supernatant was discarded and RNA extraction was performed using TRIzol™ reagent (Invitrogen) according to the manufacturer’s protocol, using equipment treated with RNaseZap™ (Invitrogen). Resultant RNA was re-suspended in 20 μL RNase-free H_2_O.

#### Tissue

10 μm sections cut from FFPE or frozen tissue blocks were stained with 1% Cresyl Violet and manually needle micro-dissected to enrich for tumour cells using a Stemi 508 Stereo microscope (Zeiss). DNA and RNA were extracted simultaneously using the AllPrep® DNA/RNA FFPE Kit (Qiagen) according to manufacturer’s instructions.

#### Blood

Anti-coagulated patient blood samples were collected where possible and processed using the DNeasy® Blood & Tissue Kit (Qiagen) according to the manufacturer’s instructions.

#### Cells

Cancer Associated Fibroblasts (CAFs) derived from the same tumours as the corresponding organoid lines were trypsinised and pelleted by centrifugation before DNA was extracted using the DNeasy® Blood and Tissue Kit (Qiagen).

### DNA sequencing

100–1500 ng of DNA per sample (ideally 1000 ng where sample permitted) was submitted to the Australian Genome Research Facility (AGRF). Whole genome sequencing (WGS) libraries were prepared using 200 Gbp xGen cfDNA & FFPE DNA Library Prep kit (IDT) with 150 paired-end reads and sequencing performed using an Illumina NovaSeq platform, aiming for >60X coverage for tumours or organoids and >30X for blood or CAF DNA (Supplementary Table 3). Sequencing reads were pre-processed with FASTP (v0.23.4)[28] and aligned using BWA-MEM2 (v2.2.1)[29] using default parameters. Point mutation variant calling, filtering and annotation was performed with SAGE (v3.4.4) and PAVE (v1.6). Copy number estimations were generated by AMBER (v4.0.1), COBALT (v1.16) and PURPLE (v4.0.2)[30]. Structural variants were called by GRIDSS2 (v2.13.2)[31] and LINX (v1.25)[32]. All these tools were run under the nf-core/oncoanalyser pipeline (v1.0.0)[33]. Tumour DNA samples were paired with their matching non-tumour DNA, when available. Variants were removed if they failed quality filters, were flagged as being low confidence or were non-pathogenic variants with an allele frequency of >0.00015 in gnomAD [34]. Whole exome sequencing (WES) libraries were prepared using Twist Biosciences Human Comprehensive Exome v2 according to the standard hybridisation protocol. One case was analysed by a targeted sequencing panel as previously described [8]. Driver genes for concordance analysis were selected from those listed in the TruSight Oncology 500 panel (Illumina, as of March 2025).

Waterfall plot and copy number (CN) frequency plots were generated in GenVisR (v 1.30.6)[35]. Overall CN segments were obtained from the somatic CN variant pipeline within Oncoanalyser (PURPLE), excluding ploidy correction and using the CN ratio compared to the matched normal sample. The CN ratio thresholds were <0.75 for loss and >1.25 for gain. Fraction of the genome altered was calculated using these segments as described previously [36]. CN segments per gene for adding *ERBB2* amplifications and *CDKN2A* deletions to the waterfall plot were absolute copy number values corrected for ploidy and purity, with thresholds >10 for amplification and <0.5 for homozygous deletion.

Organoid lines were identity profiled by the Peter Mac Callum Cancer Centre Genotyping Core Facility using short-tandem repeat (STR) loci to enable fast and reliable identification over long-term use, as well as alert users of the occurrence of any major genetic drift (Supplementary Table 4).

### RNA sequencing

50–500 ng of high-quality RNA per sample (ideally 500 ng when sufficient) was submitted to the Molecular Genomics Core, Peter MacCallum Cancer Centre for sequencing. Libraries were produced using the NEBNext Ultra II Directional Library Prep Kit with Ribo-Depletion (New England BioLabs) and sequenced on the NextSeq™ 500 System (Illumina) to obtain 75 bp single end reads. Reads were aligned to the human genome GRCh37 (hg19) using STAR [37]. Four samples had low library sizes, with two being successfully repeated. Mapped library sizes for analysed cases ranged from 3.5–26.7 M reads (average 9.6 M, Supplementary Table 3). The final sequenced cohort included three MOC cell lines (MCAS, RMUG-S, JHOM-1; two repeats each, STR verified), two normal fibroblast cell lines, five primary CAF lines, HOSE17.1 cell line (two repeats), 12 MOC tumours and 19 organoid lines (four borderline, 15 carcinoma, four with repeats).

Differential expression analysis was performed using edgeR (version 4.4.2)[38], with batch, growth pattern (expansile, infiltrative) and dose (when appropriate) in the design. Differential expression thresholds were defined as having an adjusted p-value threshold of less than 0.05 and a log fold-change threshold of at least 1.0 in either the positive or negative direction. Differentially expressed genes were visualised using pheatmap (version 1.0.12)[39] and RColorBrewer (version 1.1-3)[40].

### Single agent chemotherapy drug screens

An 80% Matrigel base layer (Corning Phenol Red Free Lot#923-100) diluted with DMEM/F-12 was prepared and 8 μL/well was dispensed to 384 well plates (384-well Flat Clear Bottom Black Corning®) using the Janus G3 Liquid Handler (Revvity). Plates were pulse centrifuged to 500 g and left to solidify at 37 °C for 20 min. Meanwhile, organoids were collected and digested to single cells using TrypLE Express (Gibco), before a manual cell count was performed using Trypan Blue and a haemocytometer. 8 µL/well of cells in a 50% Matrigel/organoid media mixture were seeded on top of the base layer to give a final density of 1500 cells/well. After a further pulse spin and incubation at 37 °C in 5% CO_2_ for 20 min to allow Matrigel/cell mixtures to set, 35 µL/well of organoid growth media was dispensed using the EL406 Liquid Handler (BioTek, Agilent). Plates were left to incubate for 72 h to allow organoids to re-form before a media change was performed with the EL406. Stock solutions of the drugs cisplatin, carboplatin, oxaliplatin, 5-fluorouracil (5-FU), gemcitabine, topotecan, irinotecan, doxorubicin, mitomycin C and staurosporine (positive control) were prepared in either DMSO or Organoid Wash Media (aqueous compounds). These were then added to cells in duplicate 10-point dilution series using the D300e Digital Dispenser (Tecan), with addition of 0.0001–0.0002% Triton-X surfactant to aqueous compounds prior to dispensing. DMSO (0.2% concentration) and Triton-X (0.0001–0.002%) were dispensed to media-containing wells as controls and concentrations standardised across drug-containing wells before plates were pulse centrifuged to remove bubbles. Plates were imaged at 4X on the Cytation C10 with a maximum projection of a stack of three z-heights as described previously [26]. Plates were then incubated for another 72 h. For plates that received a single dose, endpoint assays were then performed. Plates receiving a second dose underwent a media change, second dosing and imaging followed by incubation for 48 h before endpoint assays were performed. 10 mg/mL Hoechst 33342 (ThermoFisher Scientific) was dispensed to all wells at 1:1000 dilution using the D300e Digital Dispenser and plates were incubated for 2 hours at 37 °C. Then brightfield and fluorescent images were taken as described above using the Cytation C10. Endpoint metabolic activity was assessed by adding 25 µL/well of CellTiter-Glo (CTG, Promega) using a multichannel pipette. Plates were sealed and shaken on an orbital shaker for 20 min at room temperature before luminescence was measured at the gain of 135 on the Cytation C10.

Raw data were exported from the Cytation C10 and well images were visually assessed to identify any plate trends (e.g. uneven cell dispension, edge or location effects) and to exclude wells affected by technical artefacts (e.g. disturbed Matrigel and loss of cells). Raw CTG data was used to generate plate heatmaps and dot-plots to further assist with plate visualisation and quality control metrics. Coefficient of variation (CV) and Z’ scores were calculated across negative (DMSO and media) and positive (staurosporine 10 and 20 μM) control wells. Plates with Z’ Factor values <0 and CV values of >20% for negative controls were excluded from further analysis [41,42]. Raw CTG values of sample wells were then normalised to the median CTG of corresponding vehicle control wells, with plates divided into halves or quadrants for normalisation where significant plate-location effects were apparent. Normalised CTG values were averaged across technical replicates on each plate and used as input to the ‘auc’ function of the MESS R package v0.60 to calculate Area Under the Curve (AUC) for each drug response. Relative Area Under the Curve (RelAUC) was then calculated for each drug using the formula *RelAUC = AUC / NeutralAUC*, where *NeutralAUC= 1*.*0 × (highest concentration - lowest concentration)*. Dose-response curves for each organoid/drug combination were plotted using GraphPad Prism v10.2.3 (GraphPad Software), using the average of normalised values across biological replicate plates where available.

### Combination drug screens

Organoids were digested to single cells and plated in 384-well plates as described for single chemotherapy screens. Plates were left to incubate for 72 h to allow organoids to re-form before a media change was performed with the EL406 and drug combinations of interest (paclitaxel + carboplatin, paclitaxel + doxorubicin, paclitaxel + gemcitabine, doxorubicin + carboplatin, 5-FU + mitomycin C, 5-FU + irinotecan, 5-FU + oxaliplatin, irinotecan + oxaliplatin, and 5-FU + irinotecan + oxaliplatin) were dispensed in a 7-point dilution series at a fixed drug ratio using the Sciclone ALH 3000 Workstation. The doses were customised based on the single agent response of each individual drug in the organoids. To minimise the resource-intensive nature of high-throughput synergy screens, we employed a ‘diagonal’ matrix dose design [43]. DMSO and staurosporine controls were dispensed with the D300e Digital Dispenser and plates were pulse centrifuged. Plates were then incubated for another 72 h before endpoint assays were performed. Bright field imaging, Hoechst staining and imaging and CTG quantitation and subsequent analysis were performed as described above. Normalised CTG values for each plate were exported to the online SynergyFinder+ tool [44] that was used to predict missing response values in a full drug combination dose-response matrix. The completed matrix was used to compute synergy scores for each drug combination using the ZIP, HSA, Bliss Independence and Loewe Additivity models. Positive synergy scores were taken to indicate the presence of synergy (that the combination of drugs is more effective than the sum of the individual treatment effects), while negative synergy scores indicated antagonism.

## Results

### Growth media requirements of MOC organoids

We first attempted MOC organoid culture in 2018, before any published methods for ovarian cancer organoids were available. Extensive trials of various media formulations from colorectal and pancreatic organoid publications, in addition to testing other growth factor additives (Supplementary Figure 1), led to our first successful organoid line using MOC-ORGM3 media, uniquely formulated in our lab.

The publication of the first MOC organoids in 2019 by Kopper et al [20] detailed methods and media formulations that were similar to our own, although with some differences (Supplementary Figure 1). Subsequently, new MOC samples were trialled with both MOC- ORGM3 and Kopper media, to test if the cells of any individual tumour were better suited to growth in one or the other. Although some wells appeared to grow differently in the two media formulations at early passages, this was not consistent and generally, both media types worked equally well. RNA sequencing comparing the same organoid lines grown in different media identified only one differentially expressed gene (Supplementary Figure 1D). Therefore, after the first few passages and for all experiments described below, we used the Kopper media formulation for consistency.

As organoids derived from other cancer types are often reported to be Wnt3A dependent or independent and may in fact shift between the two according to the absence or presence of certain driver mutations [45], we set out to test this feature in a subset of diverse lines. We found that for ORG38, cells survive without Wnt3A for short periods of time, however over multiple passages of Wnt3A deprivation they began to die, suggesting an exogenous source of Wnt3A is needed for continuous growth. We expanded this experiment to six organoid lines by removing other key growth factors from the organoid medium, including each of RSPO1, Noggin, FGF10 and the combination of Wnt3A, Noggin and RSPO1. We also tested reduced Forskolin. The responses varied (Supplementary Figure 2). Reduced Forskolin was tolerated by most lines over several passages and did not affect organoid morphology. All organoid lines were universally dependent on the combination of Wnt3A/RSPO1/Noggin within 1–2 passages. RSPO1 seemed to be the single most important factor. Loss of each of Wnt3A, Noggin and FGF10 was survivable by most lines for up to three passages, but with fewer and fewer cells surviving each passage over time.

### Optimised cryopreservation methods improve MOC organoid recovery

Once the first organoid lines were established, the next challenge was cryopreservation and recovery. Finding that standard cell culture cryopreservation protocols did not reliably allow us to recover frozen organoids, we optimised our methods specifically for organoids. This included testing a range of cryoprotectant reagents, as well as the timing and extent of organoid/Matrigel manipulation prior to freezing in a robust organoid line. All freezing methods tested allowed for organoid recovery with comparable efficiencies (Supplementary Figure 3). Improved recovery was observed in instances where organoids were cryopreserved at high density rather than exponential growth phase, with the commercial Recovery Cell Culture Freezing media demonstrating the best outcome. The recovery rate was similar for digested and intact organoids, but we observed that when organoids were cryopreserved intact, they tended to recover more quickly and grow larger than if digested before freezing. For early passages or organoids that did not recover well, intact cryopreservation was therefore preferred. However, later passages and robust organoid lines were digested prior to freezing as this was more convenient for subsequent experimental use.

### Characteristics of the MOC organoid line biobank

We define a tumour organoid ‘line’ as organoids that have enduring proliferative capacity beyond three passages, can be expanded for characterisation at a genomic and phenotypic level and were able to be reliably cryopreserved and recovered. We have now successfully generated 19 MOC organoid lines (Table 1), as well as several borderline and benign mucinous ovarian tumour lines (Supplementary Table 1). Almost all MOC organoids could be continuously expanded and cultured long-term, except for ORG61, which generally struggled to continue past eight passages. The longest in our hands was cultured continuously over a year.

**Table 1:** Successful MOC organoid lines.

| ID | FIGO Stage<sup>1</sup> | Growth pattern | Grade | Sample | Morphology |
| --- | --- | --- | --- | --- | --- |
| 16 | Rec | Infiltrative | 3 | Fresh, small bowel | Dense |
| 38 | IA | Infiltrative | 3 | Fresh, ovary | Cystic/dense |
| 41 | IA | Expansile | 1 | Fresh, ovary | Cystic/dense |
| 46 | IC | Expansile | 1 | Frozen, ovary | Cystic/glandular/dense |
| 49 | IA | Infiltrative | 3 | Fresh, ovary | Cystic/glandular/dense |
| 55 | IIB | Infiltrative | 2 | Fresh, ovary | Dense |
| 59 | IC | Expansile | 1 | Fresh, ovary | Cystic bubbly |
| 60 | Rec* | Infiltrative | 3 | Fresh, ovary | Glandular/dense |
| 61 | Rec* | Infiltrative | 1 | Fresh, bowel | Cystic bubbly |
| 63 | IA | Expansile | 2 | Frozen, ovary | Glandular |
| 64 | IA | Expansile | 2 | Frozen, ovary | Glandular/cystic |
| 65 | IC | Infiltrative | 3 | Fresh, ovary | Cystic bubbly |
| 66 | IC | Expansile | 1 | Fresh, ovary | Glandular/cystic |
| 70 | IA | Expansile | 1 | Fresh, ovary | Cystic |
| 73 | IC | Expansile | 2 | Fresh, ovary | Glandular/cystic |
| 74 | IA | Expansile | 1 | Fresh, ovary | Cystic |
| 76 | Rec* | N/A | 2 | Fresh, abdominal biopsy | Dense |
| 77 | IC | Infiltrative | NA | Frozen, ovary | Glandular/dense |
| 78 | IIIC | Infiltrative | 2 | Fresh, ovary | Dense |

1. Rec = recurrent and not surgically staged at collection timepoint, \* indicates the sample we derived organoids from was the carcinoma recurrence following an initial diagnosis of mucinous borderline tumour. FIGO = International Federation of Gynaecology and Obstetrics.

The biobank includes lines derived from a range of stages and tumour grades, primary and recurrent or metastatic tumours, as well as infiltrative and expansile invasion subtypes (Figure 1B). Our success rate for generating MOC organoids after establishing optimised conditions was 70% (19/27), with failures attributable to poor tissue quality and quantity issues, including necrotic samples (n=2), cryopreserved samples (n=4), and/or samples from recurrent disease (n=3, including one with fibroblast overgrowth). Organoids were successfully generated from fresh (n=15) and viably frozen source tissue (n=4). All tumours were obtained from surgical cytoreduction or biopsy prior to any chemotherapy administration.

**Figure 1.**
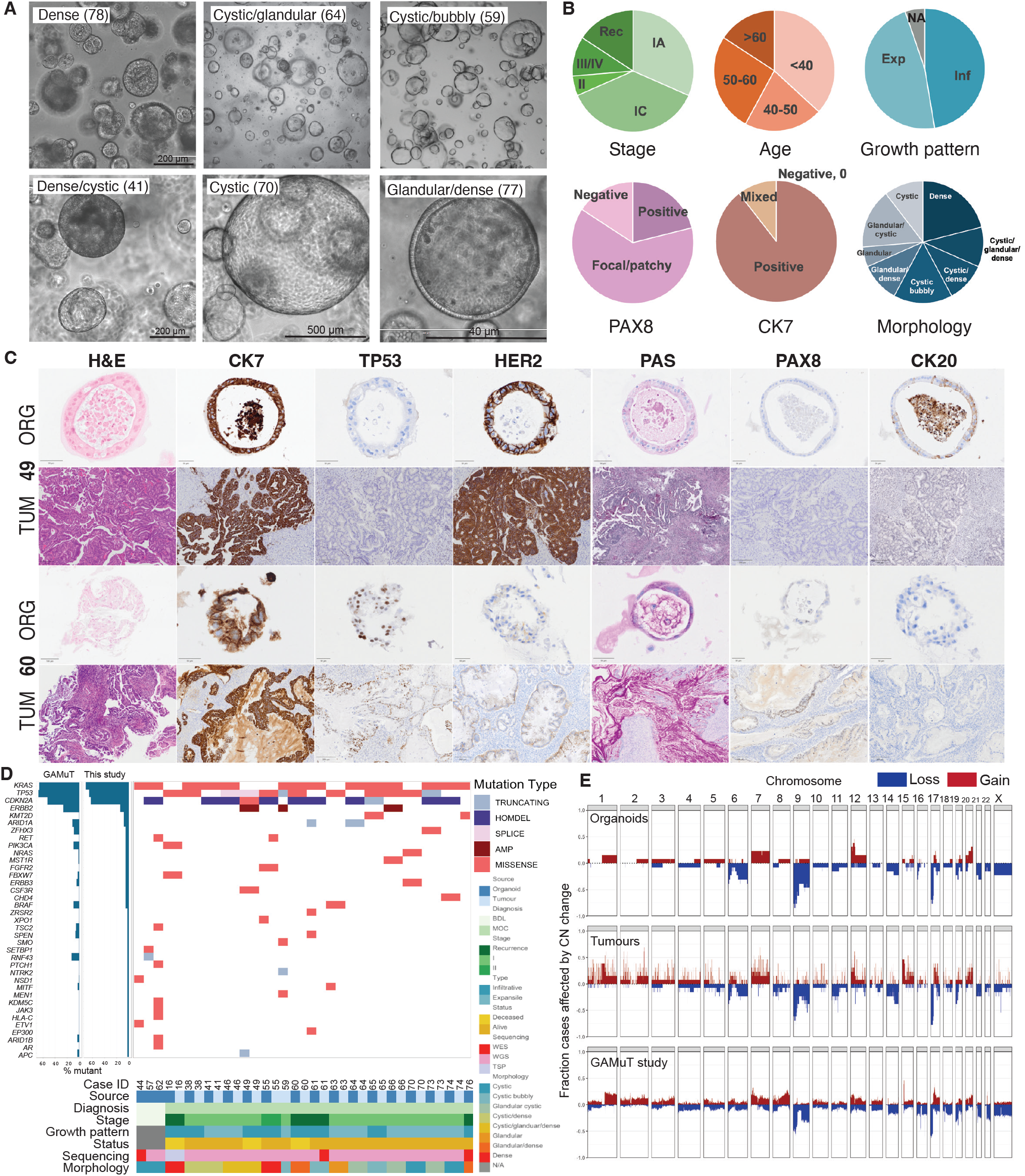
Organoid characterisation. **A.** Organoid morphology types. Cytation images (4X), and higher magnification from EVOS F1. Organoid case number in brackets. **B**. Clinico-pathological features of the primary tumours and orgnoid morphology. **C**. Immunohistochemical characterisation of two lines with their primary tumour for comparison. **D**. Plot showing somatic mutations in cancer driver genes (genes present in both Oncoanalyser and TSO500 hotspot and panel lists). Homozygous deletions (HOMDEL, CN <0.5) are only shown for *CDKN2A*. Amplifications (AMP, CN>10) are only shown for *ERBB2*. Frequency plot shown at left, with the GAMuT MOC cohort for comparison. **E**. Frequency of copy number (CN) gains (CN ratio <0.75) and losses (CN ratio >1.25) for the organoids and tumours (whole genome sequencing (WGS) data only shown) compared to the GAMuT MOC cohort. The tumours have noisier copy number because most were obtained from formalin fixed tissues. WES, whole exome sequencing; TSP, tumour sequencing panel; N/A, not available

Organoid lines showed various morphologies – some cystic with an easily distinguishable monolayer, some with multi-layered cell sheets on the outer edge but a clear lumen (“glandular cystic”), some cystic but irregular (“cystic bubbly”) and some dense with tightly packed cellular structure (Figure 1A, Supplementary Figure 4). Mixtures of morphologies were often observed within an organoid line, most commonly combinations of cystic, glandular and dense. Benign and borderline organoids were always cystic, while organoids from a carcinoma with an infiltrative growth pattern were more likely to have dense features (8/10 infiltrative with dense morphology versus 2/9 expansile, p=0.023, Fisher’s exact test). Within MOC organoids, those that contained cystic features (n=12) were more likely to be low grade (7/12 cystic were low grade, versus 0/7 not cystic, p=0.033, Fisher’s exact test) and International Federation of Gynecology and Obstetrics (FIGO) Stage I (11/12 cystic were Stage I versus 2/7 not cystic, p=0.0095, Fisher’s exact test).

IHC of diagnostic and molecular MOC markers PAS, PAX8, CK7, CK20, TP53 and ERBB2/HER2 was performed on matched organoid-tumour pairs (Figure 1C, Supplementary Table 2, Supplementary Figure 5). All but one carried a profile consistent with MOC (CK7+, CK20- or focally positive). One case included in the initial biobank was found to have high CK20 staining (ORG71) and further investigation suggested that this was a misdiagnosed gastrointestinal adenocarcinoma, possibly small bowel. This case was removed from subsequent analysis. Organoids reflected the marker expression of their parent tumours, with complete concordance in 70.5% of instances, partial concordance in 24.2% and discordance in only 5.3%. Differences most likely reflect parental tumour heterogeneity. For example, ORG60 showed focal PAX8 staining in the organoid, while the tissue was patchy positive with variable intensity of staining (Supplementary Figure 5). The least concordant organoid, ORG63, showed abnormal positive TP53 expression and negative PAX8 expression, whereas the tissue suggested wild-type staining for TP53 and positive PAX8. However, the pathology report suggested focal PAX8 staining. This tumour also carried a missense *TP53* mutation consistent with the organoid IHC staining (Figure 1D).

The least concordant marker was HER2. While 3+ and negative tumours were consistent with their organoids, tumours with 1+ or 2+ staining often had much weaker organoid staining. This observation may be due to downregulation or loss of ERBB2 expression in cases lacking gene amplification or could reflect selection *in vitro*. In the case of ORG38, for example, there was clear heterogeneity in the tumour, with areas of benign-looking epithelium having weak or no staining, borderline areas with moderate incomplete membrane staining, while carcinoma areas were negative. When combined with TP53 staining (wild-type in benign, abnormal in borderline/carcinoma, abnormal in organoids), the HER2 staining would suggest that the organoids were derived from the carcinoma areas of the original tumour.

### Organoids recapitulate parent tumour genetic profiles and greatly expand the diversity of available pre-clinical MOC models

Organoids across our biobank demonstrate the most common MOC genetic events such as *KRAS* activating mutations, homozygous deletion of *CDKN2A*, and *TP53* inactivation (Figure 1D, Supplementary Table 5). Considering all high confidence non-synonymous coding and splice site variants, the average concordance between paired tumour-organoid WGS samples was 66% (standard deviation of 23%). When variants were filtered for clinically relevant cancer driver genes, 37/44 were shared between tumour and organoid. However, after looking for the unmatched variants in Integrated Genome Viewer, three were observed to be present at either a low allele frequency or with low sequencing coverage. Therefore, the overall concordance of driver gene variants was 40/44 (91%). Across tumour-organoid pairs the average concordance was 92% (range 50–100%). Of the four variants only present in organoid (n=3) or tumour (n=1) samples, only one was predicted to be a driver mutation with high confidence. In addition to matching the variant profiles of their parent tumours, organoids also displayed similar CN variant profiles to tumours (Figure 1E, Supplementary Figure 6, Supplementary Table 6). When comparing tumours with organoids an average of 89% of each genome was concordant for CN state (gain, loss or no change) ranging from 56.6% (ORG55) to 99.9% (ORG64).

Importantly, this biobank includes models with molecular aberrations known to occur in MOC patients that are not represented in existing MOC cell lines, such as *ERBB2* amplification (ORG16, ORG49, ORG66), and mutations in *ARID1A* (ORG64), *ELF3* (ORG49), *BRAF* (ORG63) and *NRAS* (ORG70). We compared the genetic profiles represented in our biobank against the reported mutation frequencies in our previous large patient cohort and found highly similar somatic mutation and CN frequencies (Figure 1D, E).

### Transcriptomics of organoid lines

We performed RNA sequencing to compare our organoid lines with their parent tumours and other cell lines. Principal component analysis showed that samples grouped by sample type (organoid, cell line, tumour or fibroblast) rather than by case (Figure 2A). Existing MOC cell line MCAS was the most similar to the organoid lines, with RMUG-S slightly more distant, but JHOM-1 was closer to the immortalised ovarian surface epithelial cell line, HOSE 17.1, than to other MOC lines. This clustering is consistent with MCAS as a valid *in vitro* model for MOC as it carries typical MOC mutations (*KRAS, TP53*) while JHOM-1 may be more similar to an endometrioid or clear cell line (*PTEN* mutation).

**Figure 2.**
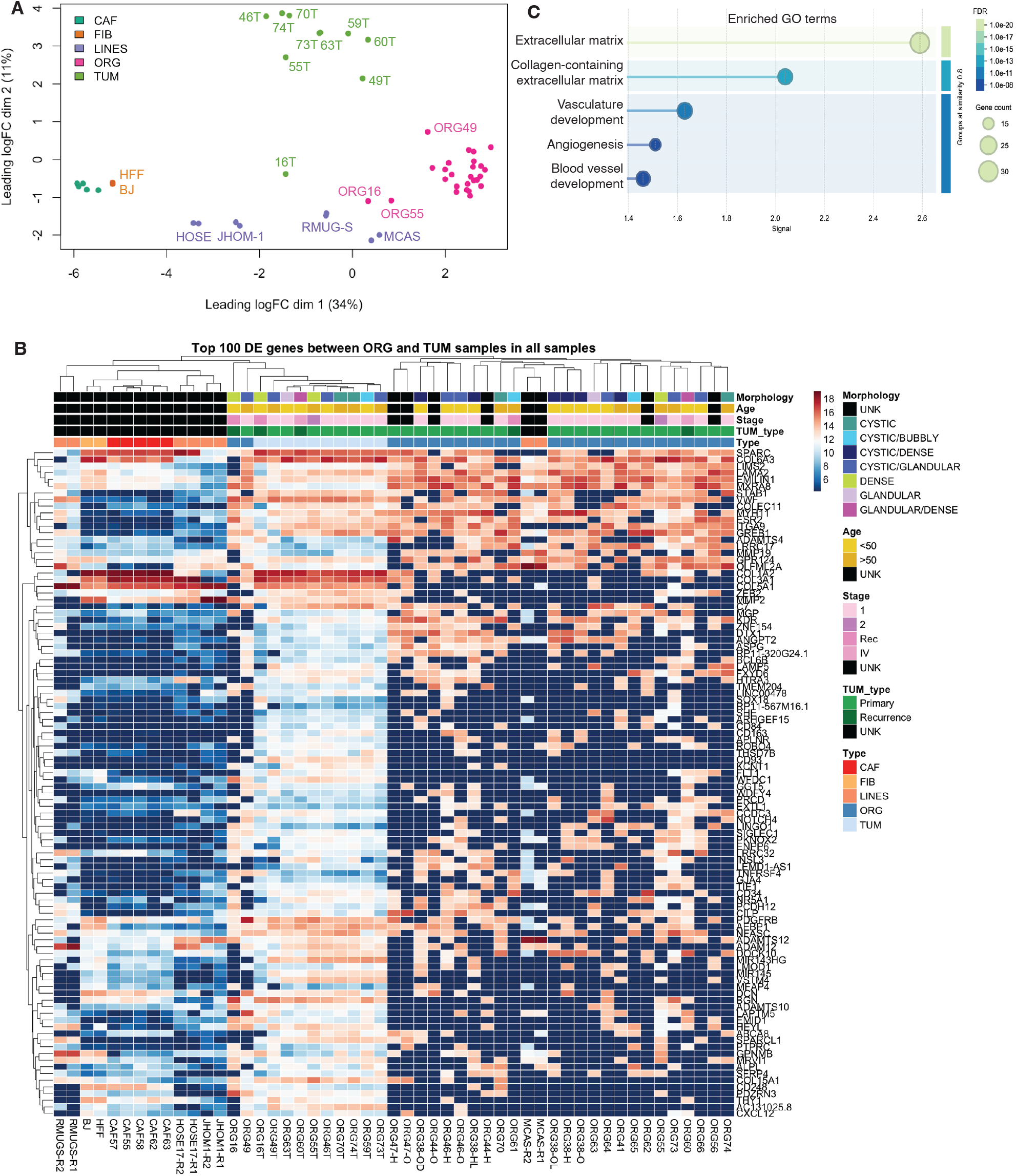
Gene expression analysis. **A.** MDS plot of all samples, coloured by cell type. CAF, cancer associated fibroblast lines; FIB - immortalised normal fibroblasts (BJ and HFF); LINES, cell lines (MCAS, RMUG, JHOM-1 and HOSE17.1), ORG, tumour organoid lines; TUM, tumour samples. **B**. Heat map showing the top 100 differentially expressed genes comparing tumours with organoid lines. **C**. Top 5 significantly enriched GO terms (by highest signal across Biological Processes, Molecular Function and Cellular Compartment using STRING (doi: 10.1093/nar/gkac1000)).

Comparing the tumours with organoids identified over 1000 differentially expressed genes (Supplementary Table 7). The tumours clustered in between the fibroblast lines and the organoids (Figure 2B). The differences between organoids and parent tumours can mostly be explained by the presence of the stromal microenvironment. Consistent with this, the top 100 differentially expressed genes were strongly enriched for genes encoding angiogenesis and extracellular matrix proteins (Figure 2C, Supplementary Table 7).

Comparing organoids to each other based on morphology found that those with a cystic morphology had downregulated genes enriched in pathways involved in multicellular organism development, the extracellular matrix (ECM) and microtubule motor activity. Dense organoids upregulated genes related to the ECM but also had higher expression of some sodium transporter genes.

### MOC organoids are sensitive to chemotherapies other than platinum agents

Given the reported platinum-resistance exhibited by MOC patient tumours, and the lack of both clinical and pre-clinical drug efficacy data specific to this ovarian cancer subtype, we used our tumour organoid lines for drug screening. We tested a panel of common chemotherapies that may be readily offered to MOC patients, chosen in consultation with treating clinicians, and compared these to the traditional ovarian standard-of-care chemotherapy drugs. Of particular interest was the potential of agents usually prescribed in gastrointestinal (GI) cancers, given the noted similarities between mucinous tumours of this origin and MOC [46]. Oxaliplatin, cisplatin, carboplatin, 5-FU, gemcitabine, mitomycin C, irinotecan, topotecan, paclitaxel, docetaxel and doxorubicin were screened across a 10-point dose dilution curve to generate dose response curves (Supplementary Figure 7), with staurosporine acting as a positive control for organoid death (Figure 3A). None of the organoid lines had been generated from parent tissues with prior exposure to cytotoxic agents. We compared adding a single chemotherapy dose (3-day drug exposure) with two doses (5-day drug exposure). Generally, the single dose 3-day drug exposure assays produced slightly reduced responses across the drug panel compared to the longer assays, (Supplementary Figure 8) but this difference was not enough to change an organoid line from being considered “resistant” to a drug to “sensitive”.

**Figure 3.**
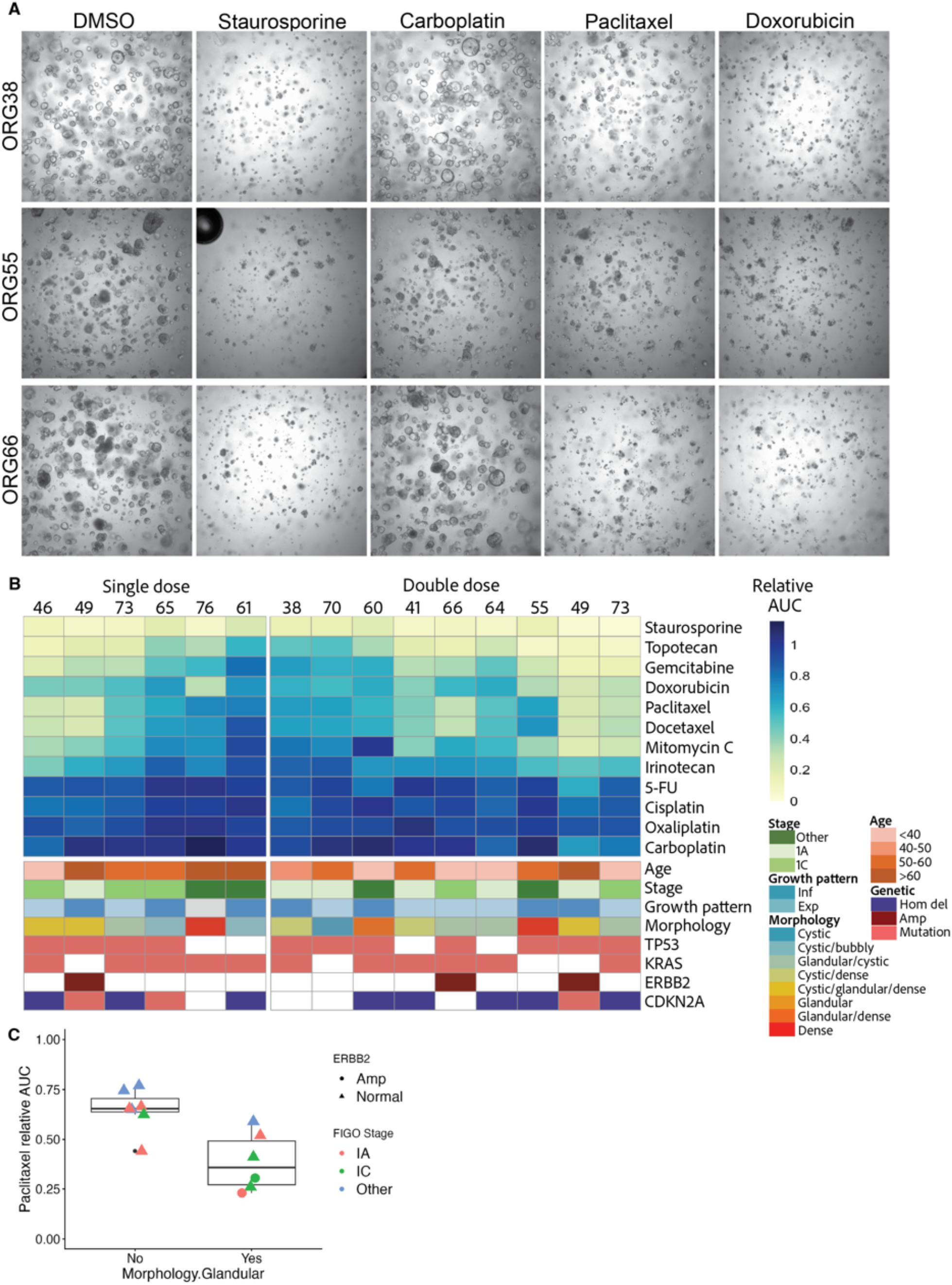
Chemotherapy screening. **A.** Brightfield images of organoids treated with DMSO or the indicated drugs at 5 µM. Staurosporine is a positive death control. **B**. Heatmap of average relative area under the curve (AUC) of CellTiterGlo (CTG) measurements from two biological repeats (each with 2 wells), except n=1 biological repeat for Organoids 46 and 60. Organoids 49 and 73 were screened with both one and two doses (one bioreplicate of each). A high relative AUC indicates resistance (blue). Drugs are ordered from most effective (top) to least, based on the average relative AUC. **C**. Relative AUC for paclitaxel by morphology and ERBB2 status, coloured by stage. Inf – infiltrative, Exp – expansile, Hom del – homozygous deletion, Amp - amplification.

Our results substantiated clinical reports of platinum resistance, with almost all organoid lines demonstrating strong resistance to all three platinum-based drugs (Figure 3B). Colorectal cancer agent 5-FU similarly had a minimal effect. Conversely, the GI cancer drug topotecan was overall the most effective agent tested, with the lowest average relative AUC across the organoid cohort.

The microtubule destabilising taxanes, paclitaxel and docetaxel, evoked similar responses in each organoid line tested. In a dose range from 0.1 µM to 1 µM these drugs consistently reduced cell viability by 25–75%, only showing a dose response at very low doses (Supplementary Figure 9). As the concentration increased, viability as measured by CTG did not decrease further, and indeed appeared to increase at the highest doses (5–10 µM) (Supplementary Figure 7).

A number of agents including mitomycin C, doxorubicin, and the topoisomerase I inhibitor topotecan showed good efficacy particularly at high concentrations, with typical dose-response curves produced. Irinotecan, another topoisomerase I inhibitor, was also effective in some lines, with increased cytotoxicity observed when dosing with the active component of the drug, SN38. Mitomycin C generated the most diverse response across the organoid lines tested, with insensitive (e.g. ORG38, ORG60) and sensitive lines (e.g. ORG49, ORG73).

To investigate differences in individual organoid line drug sensitivity profiles, subgroup analysis was performed, dividing organoids according to genetic profiles, stage, grade, age, tumour growth pattern and organoid morphology. There was no statistically significant association between organoid drug response and tumour growth pattern, patient age, *KRAS* or *TP53* mutation, fraction of the genome altered by copy number, ploidy or whole genome duplication, either by dichotomising sensitivity or using the relative AUC values in non-parametric t-tests, ANOVA tests or Spearman correlations. Although associations were not statistically significant once multiple testing correction was performed, there was a trend for organoid sensitivity to the taxanes paclitaxel and docetaxel to be associated with FIGO Stage I tumours (Wilcoxon t-test p=0.05 and p=0.02 respectively, uncorrected) and *ERBB2* amplification (Wilcoxon t-test p=0.051 for both). Both of the evaluated organoid lines with *ERBB2* amplification were among the most sensitive to paclitaxel (Figure 3C). A better paclitaxel response was also associated with glandular organoid morphology (Wilcoxon t-test p=0.005, uncorrected). A model incorporating all three aspects (stage, *ERBB2* and glandular morphology) had the best fit for paclitaxel response compared to any one or two of these factors (Akaike Information Criterion value –19.2 versus –14.6 for next lowest, which was *ERBB2* with morphology).

We then evaluated the baseline gene expression profiles of 12 organoid lines with respect to their drug responses. Firstly, we tested for genes that were correlated with drug response by using a generalised linear model and modelling each organoid with its corresponding AUC on a continuous scale for each drug. We only tested those drugs with sufficiently varying responses (standard deviation of relAUC >0.1). After correction for false discovery, this approach only identified five statistically significantly differentially expressed genes that were correlated with mitomycin C response (*IL34, TSHZ2, ATHL1, RP11-326C3,2* and *MAP7D3*). We performed a second differential expression test between organoids by ranking organoid lines by AUC and classifying each as “sensitive” (relative AUC <= 0.5 or “resistant” (relative AUC >= 0.7). Only mitomycin C had enough cases (at least three) in each group for an analysis, which identified just one differentially expressed gene (*SAA1*). In addition, we considered the average relative AUCs for drug classes, for which cytotoxic antibiotics (doxorubicin and mitomycin C) was the only class to have enough cases for analysis. Three differentially expressed genes were detected for this class: *SAA1, TSHZ2* and *TFF1*.

### Synergy screening of MOC organoid lines

Acknowledging that most of the chemotherapies tested on our organoids are commonly prescribed as part of combination rather than single agent regimens, we next performed combination screens in six organoid lines representing a range of single agent responses. We tested the standard ovarian cancer regimen carboplatin + paclitaxel and the alternative GI cancer regimens 5-FU + oxaliplatin (FOLFOX), 5-FU + irinotecan (FOLFIRI) and 5-FU + oxaliplatin + irinotecan (FOLFOXIRI). This screen replicated the results of the single agent screen for individual chemotherapy dose response curves (Supplementary Figure 9). Carboplatin + paclitaxel showed at least an additive effect in ORG64 and ORG60 (Figure 4A, B, C) but was not significantly different compared to paclitaxel alone across all lines. 5-FU + oxaliplatin showed no synergistic effect, however, across all lines 5-FU + irinotecan and 5-FU + oxaliplatin + irinotecan had significantly lower AUC values compared to the respective “anchors” (i.e. the common drug to which others were added, Figure 4A). However, this difference was not reflected in the synergy scores for individual lines, which generally showed antagonism for 5-FU combinations (Figure 4B). This result reflects that 5-FU is not effective alone, and the lower AUC was due to the effect of irinotecan. When a paired t-test was performed for irinotecan alone versus 5-FU + irinotecan or 5-FU + oxaliplatin + irinotecan there was no significant difference (p=0.8 and 0.46 respectively).

**Figure 4.**
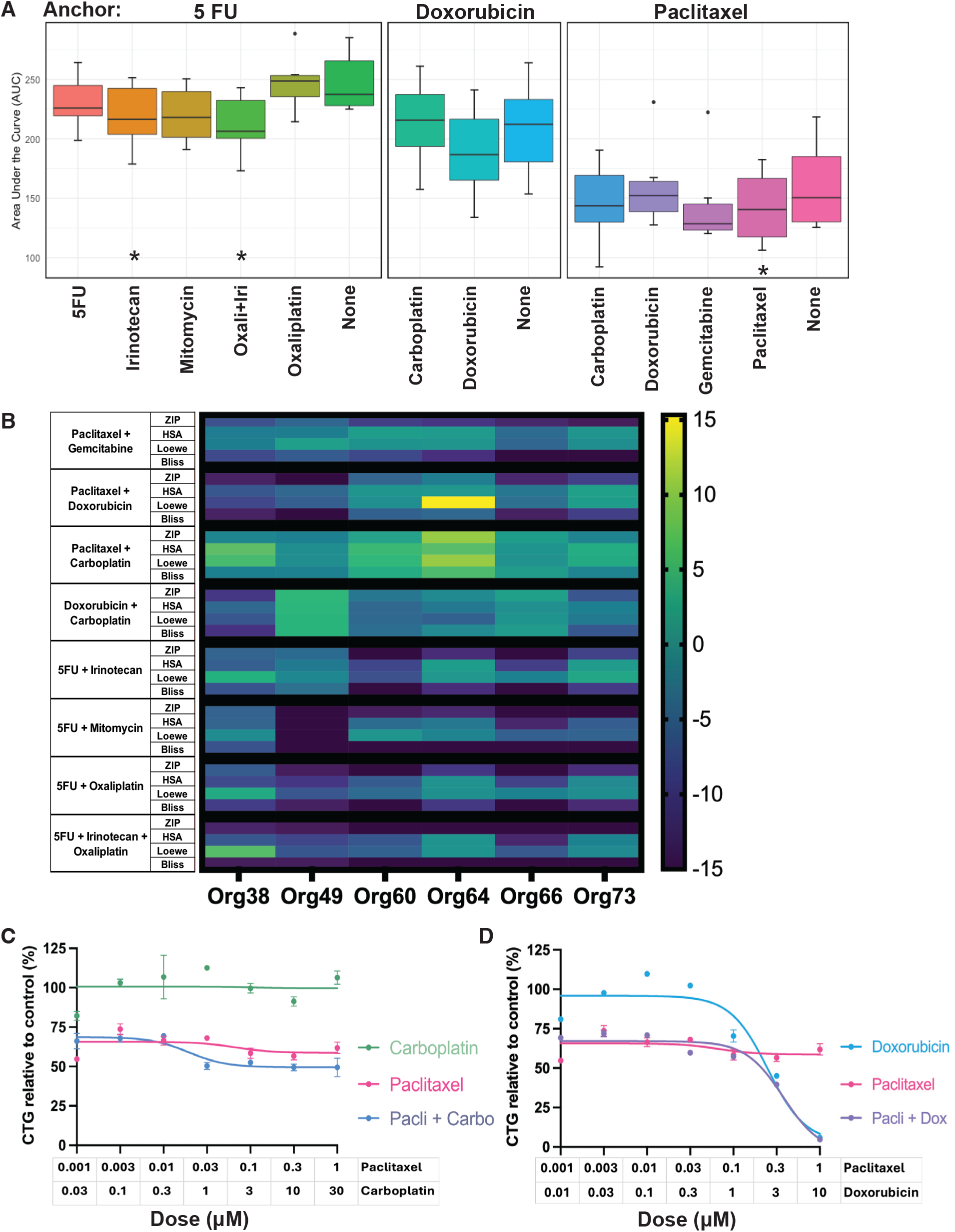
Drug combination screening. **A.** For each anchor, the area under the curve (AUC) values are shown for the anchor and its combinations. Plots are the results for six organoid lines with median, inter-quartile range and 95% confidence interval shown. * p<0.05; paired t-test comparison to the anchor with false discovery rate correction. **B**. Synergy scores across all combinations in six organoid lines and four methods of calculating synergy. 0, neither additive nor antagonistic; >0, additive or synergistic; <0 antagonistic. **C**. Dose response for ORG64 comparing paclitaxel with carboplatin and the combination. **D**. Dose response for ORG64 comparing paclitaxel with doxorubicin and the combination.

We also investigated less-usual combinations that were of interest to clinicians and involved the most active chemotherapies from our single agent screens, including doxorubicin and gemcitabine. No consistent synergistic combinations were observed (Figure 4A) however ORG64 showed synergy between paclitaxel + doxorubicin (Figure 4D), while ORG49 showed at least additive effects of carboplatin + doxorubicin.

### Case study: Organoid screening of advanced MOC from a percutaneous biopsy

Patient 76 (corresponding to ORG76) was aged in her 60s when she was found to have lung and peritoneal metastases three years after an initial diagnosis of a non-invasive mucinous borderline ovarian tumour. We received three radiologically-guided biopsy cores from a peritoneal nodule that were processed for organoids. Within three months, organoids were successfully grown and a screen performed using 11 chemotherapy agents (Figure 5). Analysis showed that relative to other patient organoids, this tumour was sensitive to doxorubicin, somewhat resistant to carboplatin and had an average response to paclitaxel. By the time the results were returned, the patient had been treated with carboplatin and paclitaxel, which was effective initially, but disease progression occurred within four months of therapy completion. As there are no standard-of-care guidelines for second-line therapy and the patient was deteriorating and too unwell for clinical trials, the treating clinician elected to trial doxorubicin based on the results of the organoid drug screen. The patient showed an excellent response to this treatment, with substantial improvement in symptoms, performance status and quality of life and a good partial response on imaging. However, after completing the treatment course and transitioning from doxorubicin to maintenance bevacizumab, she again developed symptoms. She enrolled into a Phase I clinical trial that aimed to target the *KRAS* mutation in the tumour, however experienced complications from treatment and had to withdraw from the trial, passing away some months later.

**Figure 5.**
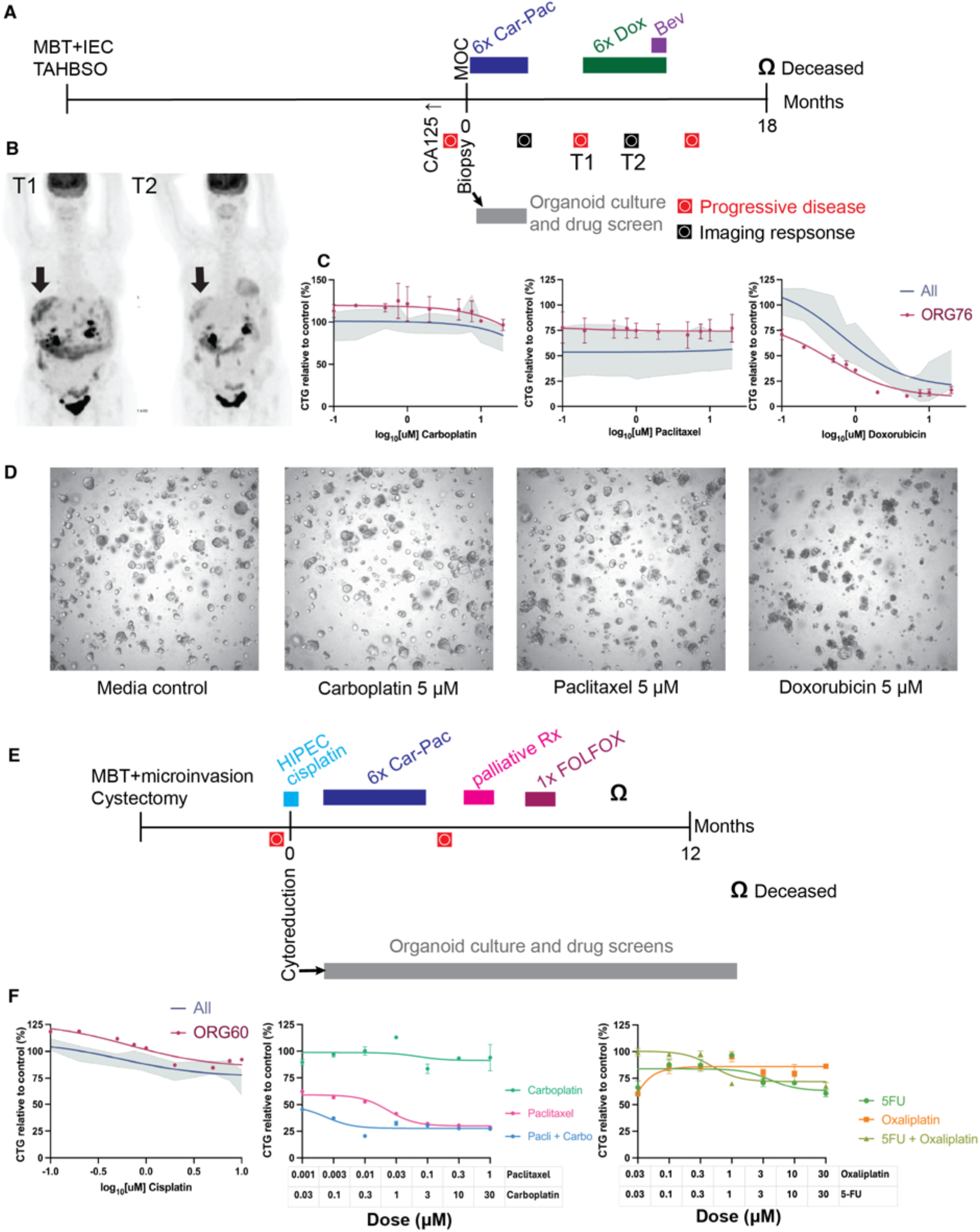
Clinical case studies. **A.** Timeline for Patient 76 (not to scale). MBT+IEC - mucinous borderline tumour with intra-epithelial carcinoma. TAHBSO - total hysterectomy and bilateral salpingo-oophorectomy. Car-Pac - carboplatin-paclitaxel. Dox - doxorubicin. T1, T2 – first and second imaging timepoints shown in **B**. FDG/PET CT imaging. **C**. Dose response curves comparing ORG76 (two bioreplicates, each the average of duplicate wells) with the average and 95% confidence interval of all other samples with single dose treatments (1–2 biological repeats, each the average of duplicate wells). **D**. Images of ORG76 treated with the indicated drugs. **E**. Patient 60 timeline. HIPEC - heated intra- peritoneal chemotherapy. FOLFOX - 5-FU, oxaliplatin, leucovorin (incomplete). **F**. Dose response curves of single agent cisplatin, comparing ORG60 to the other organoid lines with two dose treatments, as well as responses to the used combination regimens carboplatin-paclitaxel and FOLFOX.

After this case, we investigated the feasibility of organoid drug screening in a clinically relevant timeframe. We tested tissue received from one case with advanced stage MOC (ORG78) and a case of recurrent MOC for which we had cryopreserved tissue from the primary tumour but had not made organoids at the time of diagnosis (ORG77). Organoids were established from both cases within two weeks, and a first drug screen using a reduced plate layout and single dosing with drug-specific dose ranges was performed within four weeks (Supplementary Figure 10). The results for these cases were consistent with previous organoid lines.

### Further correlations with clinical outcomes

The majority of patients from which the tumour organoids were derived were Stage I, and as such only two, patient 76 as described above and patient 55 were offered chemotherapy. The latter patient had Stage IIB MOC, with complete cytoreduction and adjuvant carboplatin/paclitaxel. She remained progression-free with three years of follow-up. ORG55 was classed as sensitive overall, but relatively resistant to both carboplatin and paclitaxel. Patient 49 did not have any adjuvant chemotherapy but recurred with lung metastasis after 20 months. She received carboplatin/paclitaxel but died within a month of her recurrence being detected.

Patient 60 recurred within nine months of a cystectomy for a borderline mucinous tumour. Her extensive disease was unable to be completely resected and she underwent heated intra-peritoneal chemotherapy (HIPEC) using cisplatin, followed by carboplatin/paclitaxel. Her disease progressed within six months. Switching to FOLFOX was ineffective and she died 10 months after the HIPEC procedure. ORG60 was classed as overall resistant to chemotherapy, although the cells did show an additive effect of the combination of carboplatin and paclitaxel. The combination of 5-FU and oxaliplatin was not additive and under some models was classed as antagonistic.

Patient 61 was highly unusual, having survived a high-grade serous ovarian carcinoma 18 years prior to her mucinous tumour diagnosis. She had multiple borderline lesions seven and six years before recurring with extensive adenocarcinoma. Her complete cytoreductive surgery for this tumour was followed by carboplatin, and she was progression-free with 31 months of follow-up. ORG61 was the most resistant of any of the organoid lines overall, but the most sensitive to carboplatin. The long timeframe of progression, the difficulty in maintaining long-term culture of this line and the quiet genomic profile (*KRAS* mutation, *CDKN2A* deletion but no other CN events) suggest that this tumour could be somewhat indolent clinically.

## Discussion

Here we present the world’s largest and most comprehensive mucinous ovarian tumour organoid cohort. This represents a major advance in the study of MOC, as the historical lack of pre-clinical models specific to this histotype has greatly impeded progress. This organoid biobank captures a range of clinico-pathological features and somatic mutation profiles to greater reflect the molecular heterogeneity seen in the patient population, significantly expanding upon the diversity of the few pre-existing MOC *in vitro* models. With the dearth of clinical-trial informed data we posit this cohort of MOC models will serve as an invaluable vehicle to explore more effective therapies for patients with MOC.

While the development of a method to generate and culture MOC organoids was initially challenging, our protocols are now reliable and consistently successful, with a 70% success rate of line generation across fresh, frozen and biopsy samples collected. This rate is similar to others in the literature for ovarian cancer that range from 44–65%, although HGSOC generally has lower rates of success [25,47-51]. Most lines retain proliferative capacity over many passages, like Kopper *et al*, and in contrast to Maenhoudt *et al*, whose MOC organoid line was not able to be maintained long-term perhaps due to the lack of Wnt3A in their medium. The importance of Wnt3A for MOC organoid growth contrasts with reports that HGSOC organoids require a low-Wnt3A, and perhaps also a low RSPO1 environment for growth [47,50], highlighting potential inherent biological differences between these ovarian cancer histotypes. This is noteworthy in light of previous studies suggesting Wnt-pathway inhibitors should be investigated as a potential targeted therapy for MOC, given that 8–12% of MOC tumours have been shown to harbour inactivating mutations in *RNF43*, a known negative inhibitor of Wnt signalling [8]. However, only ORG57, derived from a borderline tumour, carried an inactivating *RNF43* mutation in our cohort. The sensitivity to media components related to the Wnt pathway may therefore be intrinsic to MOC and not determined by genetic events.

The MOC organoids largely recapitulate parent tumour characteristics as indicated by genetic and IHC analysis. Overall, truncal driver mutations were highly concordant, however CN events were more variable. Where organoids did not completely match parent tumours, we suspect this is due to heterogeneity in the tumour sections used for DNA extraction, such as incorporating both borderline and invasive tumour cells, as opposed to the organoids. While selection may occur in culture with one lineage coming to dominate the population at time of sequencing, the passage used for sequencing for paired samples was quite early (median passage number = 3).

In performing high-throughput drug screens using our organoid cohort, we present the first extensive pre-clinical drug efficacy dataset for this rare cancer, both for single agents and combinations. We were able to achieve this important milestone in the field by optimising culture conditions and miniaturising organoid growth to a 384-well format for screening. Clinical trial data for MOC is difficult to obtain due to the numbers of participants required to achieve statistical power and thus there has historically been little evidence available to guide treating clinicians.

Importantly, the organoid lines showed consistent and limited responses to all platinum agents, with none showing any response below 20 μM, whereas HGSOC organoids from platinum-sensitive patients had an IC50 of 5–10 μM [52]. While most of our drug assays utilised a 20 μM maximum dose, more recent assays incorporating higher platinum drug doses up to 150 μM still demonstrated minimal MOC organoid responses (Supplementary Figure 10). Our findings of platinum resistance are concordant with clinical reports and may account for poor survival of late-stage MOC patients, who are most often treated with first line carboplatin/paclitaxel. Similarly, although all lines respond somewhat to paclitaxel, they show a plateau in their response curves, suggesting a remnant population of cells that survive treatment. This plateau effect, which ranged from 25–75% of the DMSO control, remained in the combination treatments, even in the organoid lines with at least an additive effect of the two drugs in combination. Adding gemcitabine or doxorubicin to paclitaxel did have an additive effect in most organoid lines, particularly doxorubicin, but in 5/6 a stubborn proportion of cells (approximately 15–30%) remained.

The organoids also showed a limited response to drugs that comprise a common GI tumour regimen, FOLFOX/CapOX. Although an earlier *in vitro* study had suggested this as a potential treatment [53], the mEOC/GOG241 clinical trial did not show a significant difference in response to CapOx vs carboplatin + paclitaxel, though the sample size was small [54]. Retrospective data studies have seen conflicting signals for GI versus ovarian regimens, but it is possible these could be confounded by year of diagnosis, clinico-pathological features such as tumour grade, patient age and ECOG status, and treatment intent [55,56]. Hence, larger prospective studies are needed to be able to account for these issues. However, the *in vitro* data described here could not identify a strong signal to support the use of a GI regimen containing 5-FU or capecitabine preferentially to an ovarian regimen.

Several alternate chemotherapies, all well-established clinically in the treatment of other cancers, showed high efficacy in MOC organoids. Based on these results, gemcitabine, doxorubicin, mitomycin C and topotecan could be considered for use in the treatment of MOC patients. Others have reported the efficacy of irinotecan, mitomycin C and doxorubicin over cisplatin [57], although this work was conducted on only one cell line, OMC-3, that has since been deemed as unlikely to represent primary MOC due to its mutational profile [7]. A small phase II clinical trial previously demonstrated increased complete/partial response to an irinotecan/docetaxel over carboplatin/paclitaxel regimen, however only four MOC patients were included, and there was found to be no difference in overall or progression-free survival [58]. Future *in vivo* studies using *in vitro* screens to build further evidence for the use of these agents and provide clinicians with confidence to trial their patients on these therapies.

We did not find robust correlates of chemotherapy response at an RNA level. The absence of these correlates may reflect the heterogeneity of chemotherapy resistance mechanisms as well as the limited power of our analysis. Our data do not support previous work describing oxaliplatin resistance due to PRKRA/PACT expression [4], with no statistically significant correlation between *PRKCA* RNA expression and the AUC values calculated for oxaliplatin. Although others have found differences in expression using similarly powered RNAseq and organoid AUC in HGSOC [59], the paucity of findings here may reflect a limited dynamic range of responses for some drugs (e.g. carboplatin) and/or more diverse mechanisms of resistance.

The association of *ERBB2* amplification with paclitaxel response is interesting but only based on two cases. Nonetheless, a correlation between *ERBB2* amplification or overexpression and paclitaxel response has been observed in multiple studies of breast cancer [60-62]. Given that *ERBB2* amplification is associated with the expansile growth pattern and with improved outcomes in MOC [63], it will be challenging to find similar clinical data to determine the broader significance.

With the current success ratescreen drugs prior to use in that patient, within a clinically relevant timeframe. Our experience suggests that this is most easily performed when ample surgical tissue is received (as for the screen performed within 2 weeks for ORG78), but we have demonstrated that this is feasible even with low volume tissue collected from percutaneous biopsy. Drug treatments of organoid models are reported to show good correlation with patient outcomes [13]. A limitation of this study is that it was not prospective and response assessments were inferred retrospectively from clinical records rather than being evaluated in real time using standardized criteria such as RECIST. In addition, most patients did not receive systemic chemotherapy, and for those that did, complete cytoreduction meant that it was difficult to determine if the reason a patient did well was attributable to response of the tumour cells to chemotherapy. Nonetheless, it was reassuring that a patient whose organoid was sensitive to doxorubicin also received substantial clinical benefit while on treatment with this drug. The case study reported here also suggests that organoids grown from tissue obtained before therapy can still reflect patient response even in the relapsed setting and may inform second-line therapy selection.

In addition to their utility for pre-clinical drug screening, including relevant targeted therapies, the organoids are also amenable to a range of other research applications. However, these models remain limited compared to their *in vivo* counterparts, and the next challenge is to reach higher level tissue complexity, incorporating more differentiated cell types and other aspects of the microenvironment, to more closely mimic heterotypic biological systems.

## Acknowledgements

This study was supported by the Congressionally Directed Medical Research Program of the US Dept of Defense (OC170121 and OC200056), The National Health and Medical Research Council Australia (NHMRC Investigator funding to: AO, GNT1196256 and GNT2033499; CLS, GNT2009783; ADeF, GNT2033042), The Peter MacCallum Foundation, and The Stafford-Fox Rare Cancer Program. CS, ND and OC were supported by the University of Melbourne Postgraduate Research Scholarships. We thank the Peter MacCallum Cancer Centre Genotyping Core (RRID:SCR_025622), Centre for Advanced Histology and Microscopy (RRID:SCR_025432), Victorian Centre for Functional Genomics (RRID:SCR_025582), Molecular Genomics Core (RRID:SCR_025695), Research Laboratory Support Services (RRID:SCR_025699) and Bioinformatics Core (RRID: SCR_025901). These facilities are supported by the Peter MacCallum Foundation, the Australian Cancer Research Foundation, the Victorian State Government Operational Infrastructure Support and Australian Government NHMRC IRIISS. We thank the Victorian Cancer Biobank for sample collection and Prof Sean Grimmond (University of Melbourne) for assistance with DNA sequencing. We thank Tim Semple, Hugo Saunders, Xiang Mark Li and Niko Thio for assistance with sequencing and analysis pipelines. The Victorian Centre for Functional Genomics (K.J.S) is funded by Phenomics Australia (https://ror.org/0201hm243), through funding from the Australian Government’s National Collaborative Research Infrastructure Strategy (NCRIS) program and the University of Melbourne Collaborative Research Infrastructure Program. The Gynaecological Oncology Biobank at Westmead was funded by the NHMRC (ID310670, ID628903); the Cancer Institute NSW (12/RIG/1-17, 15/RIG/1-16); the Department of Gynaecological Oncology, Westmead Hospital; and acknowledges financial support from the Sydney West Translational Cancer Research Centre, funded by the Cancer Institute NSW (15/TRC/1-01). This research was supported by NSW Health Pathology through its contribution of patient samples and pathology data. We thank Nadia Traficante and Leanne Bowes from the Australian Ovarian Cancer Study (AOCS) for assistance with patient recruitment. AOCS was supported by the U.S. Army Medical Research and Materiel Command under DAMD17-01-1-0729, The Cancer Council Victoria, Ovarian Cancer Australia and the Peter MacCallum Foundation. We gratefully acknowledge and sincerely thank the patients and families who participated in this study. Their generous contributions made this work possible.

## Notes

### Competing Interest Statement

The authors have declared no competing interest.

## References

1. Peres LC, Cushing-Haugen KL, Kobel M, et al. Invasive Epithelial Ovarian Cancer Survival by Histotype and Disease Stage. J Natl Cancer Inst 2019; 111: 60–68.

2. Zaino RJ, Brady MF, Lele SM, et al. Advanced stage mucinous adenocarcinoma of the ovary is both rare and highly lethal: a Gynecologic Oncology Group study. Cancer 2011; 117: 554–562.

3. Affatato R, Carrassa L, Chilà R, et al. Identification of PLK1 as a New Therapeutic Target in Mucinous Ovarian Carcinoma. Cancers 2020; 12: 672.

4. Hisamatsu T, McGuire M, Wu SY, et al. PRKRA/PACT Expression Promotes Chemoresistance of Mucinous Ovarian Cancer. Mol Cancer Ther 2019; 18: 162–172.

5. Kidera Y, Yoshimura T, Ohkuma Y, et al. [Establishment and characterization of a cell line derived from mucinous cystadenocarcinoma of human ovary]. Nihon Sanka Fujinka Gakkai Zasshi 1985; 37: 1820–1824.

6. Sakayori M, Nozawa S, Udagawa Y, et al. [Biological properties of two newly established cell lines (RMUG-S, RMUG-L) from a human ovarian mucinous cystadenocarcinoma]. Hum Cell 1990; 3: 52–56.

7. Craig O, Nigam A, Dall GV, et al. Rare Epithelial Ovarian Cancers: Low Grade Serous and Mucinous Carcinomas. Cold Spring Harb Perspect Med 2023; 13.

8. Cheasley D, Wakefield MJ, Ryland GL, et al. The molecular origin and taxonomy of mucinous ovarian carcinoma. Nat Commun 2019; 10: 3935.

9. Alexandre J, Ray-Coquard I, Selle F, et al. Mucinous advanced epithelial ovarian carcinoma: clinical presentation and sensitivity to platinum-paclitaxel-based chemotherapy, the GINECO experience. Ann Oncol 2010; 21: 2377–2381.

10. Bamias A, Psaltopoulou T, Sotiropoulou M, et al. Mucinous but not clear cell histology is associated with inferior survival in patients with advanced stage ovarian carcinoma treated with platinum-paclitaxel chemotherapy. Cancer 2010; 116: 1462–1468.

11. Shimada M, Kigawa J, Ohishi Y, et al. Clinicopathological characteristics of mucinous adenocarcinoma of the ovary. Gynecol Oncol 2009; 113: 331–334.

12. Hollis RL, Stillie LJ, Hopkins S, et al. Clinicopathological Determinants of Recurrence Risk and Survival in Mucinous Ovarian Carcinoma. Cancers 2021; 13: 5839.

13. Vlachogiannis G, Hedayat S, Vatsiou A, et al. Patient-derived organoids model treatment response of metastatic gastrointestinal cancers. Science 2018; 359: 920–926.

14. Weeber F, van de Wetering M, Hoogstraat M, et al. Preserved genetic diversity in organoids cultured from biopsies of human colorectal cancer metastases. Proceedings of the National Academy of Sciences of the United States of America 2015; 112: 13308–13311.

15. Kollmann C, Buerkert H, Meir M, et al. Human organoids are superior to cell culture models for intestinal barrier research. Front Cell Dev Biol 2023; 11: 1223032.

16. Sun T, Jackson S, Haycock JW, et al. Culture of skin cells in 3D rather than 2D improves their ability to survive exposure to cytotoxic agents. J Biotechnol 2006; 122: 372–381.

17. Fatehullah A, Tan SH, Barker N. Organoids as an in vitro model of human development and disease. Nat Cell Biol 2016; 18: 246–254.

18. Sato T, Vries RG, Snippert HJ, et al. Single Lgr5 stem cells build crypt-villus structures in vitro without a mesenchymal niche. Nature 2009; 459: 262–265.

19. Drost J, Clevers H. Organoids in cancer research. Nature Reviews Cancer 2018; 18: 407–418.

20. Kopper O, de Witte CJ, Lõhmussaar K, et al. An organoid platform for ovarian cancer captures intra- and interpatient heterogeneity. Nature Medicine 2019; 25: 838–849.

21. de Witte CJ, Espejo Valle-Inclan J, Hami N, et al. Patient-Derived Ovarian Cancer Organoids Mimic Clinical Response and Exhibit Heterogeneous Inter- and Intrapatient Drug Responses. Cell Reports 2020; 31: 107762.

22. Maenhoudt N, Defraye C, Boretto M, et al. Developing Organoids from Ovarian Cancer as Experimental and Preclinical Models. Stem Cell Reports 2020; 14: 717–729.

23. Kim J, Seo JH, Kim Y-H, et al. Abstract 6042: Developing patient-derived organoids and tumor xenograft model for ovarian cancer for preclinical therapeutic evaluation. Cancer Research 2022; 82: 6042–6042.

24. Ledermann JA, Luvero D, Shafer A, et al. Gynecologic Cancer InterGroup (GCIG) consensus review for mucinous ovarian carcinoma. Int J Gynecol Cancer 2014; 24: S14–19.

25. Kopper O, de Witte CJ, Lohmussaar K, et al. An organoid platform for ovarian cancer captures intra- and interpatient heterogeneity. Nat Med 2019; 25: 838–849.

26. Choo N, Ramm S, Luu J, et al. High-Throughput Imaging Assay for Drug Screening of 3D Prostate Cancer Organoids. SLAS Discov 2021; 26: 1107–1124.

27. Kang EY, Cheasley D, LePage C, et al. Refined cut-off for TP53 immunohistochemistry improves prediction of TP53 mutation status in ovarian mucinous tumors: implications for outcome analyses. Mod Pathol 2021; 34: 194–206.

28. Chen S, Zhou Y, Chen Y, et al. fastp: an ultra-fast all-in-one FASTQ preprocessor. Bioinformatics 2018; 34: i884–i890.

29. Vasimuddin M, Misra S, Li H, et al. Efficient Architecture-Aware Acceleration of BWA-MEM for Multicore Systems. In: 2019 IEEE International Parallel and Distributed Processing Symposium (IPDPS). (ed)^(eds), 2019; 314–324.

30. Priestley P, Baber J, Lolkema MP, et al. Pan-cancer whole-genome analyses of metastatic solid tumours. Nature 2019; 575: 210–216.

31. Cameron DL, Baber J, Shale C, et al. GRIDSS2: comprehensive characterisation of somatic structural variation using single breakend variants and structural variant phasing. Genome Biology 2021; 22: 202.

32. Shale C, Cameron DL, Baber J, et al. Unscrambling cancer genomes via integrated analysis of structural variation and copy number. Cell Genomics 2022; 2.

33. Stephen Watts MC, Luan Nguyen, nf-core bot, charlesshale, Cassie Litchfield, Matthijs van Niekerk, Simone Coughlan, Ömer An, Simon Pearce, & Rayan Hassaïne. nf-core/oncoanalyser. In. Pied Currawong 1.0.0 ed. (ed)^(eds). Zenodo, 2024.

34. Karczewski KJ, Francioli LC, Tiao G, et al. The mutational constraint spectrum quantified from variation in 141,456 humans. Nature 2020; 581: 434–443.

35. Skidmore ZL, Wagner AH, Lesurf R, et al. GenVisR: Genomic Visualizations in R. Bioinformatics 2016; 32: 3012–3014.

36. Cheasley D, Wakefield MJ, Ryland GL, et al. The molecular origin and taxonomy of mucinous ovarian carcinoma. Nature Communications 2019; 10: 3935.

37. Dobin A, Davis CA, Schlesinger F, et al. STAR: ultrafast universal RNA-seq aligner. Bioinformatics 2013; 29: 15–21.

38. McCarthy DJ, Chen Y, Smyth GK. Differential expression analysis of multifactor RNA-Seq experiments with respect to biological variation. Nucleic Acids Res 2012.

39. Kolde R. Pheatmap: pretty heatmaps. R package version 2019; 1: 726.

40. Neuwirth E. RColorBrewer: colorbrewer palettes.

41. Birmingham A, Selfors LM, Forster T, et al. Statistical methods for analysis of high-throughput RNA interference screens. Nat Methods 2009; 6: 569–575.

42. Zhang JH, Chung TD, Oldenburg KR. A Simple Statistical Parameter for Use in Evaluation and Validation of High Throughput Screening Assays. J Biomol Screen 1999; 4: 67–73.

43. Ianevski A, Giri AK, Gautam P, et al. Prediction of drug combination effects with a minimal set of experiments. Nat Mach Intell 2019; 1: 568–577.

44. Zheng S, Wang W, Aldahdooh J, et al. SynergyFinder Plus: Toward Better Interpretation and Annotation of Drug Combination Screening Datasets. Genomics Proteomics Bioinformatics 2022; 20: 587–596.

45. Han T, Goswami S, Hu Y, et al. Lineage Reversion Drives WNT Independence in Intestinal Cancer. Cancer Discov 2020; 10: 1590–1609.

46. Kelemen LE, Kobel M. Mucinous carcinomas of the ovary and colorectum: different organ, same dilemma. The lancet oncology 2011; 12: 1071–1080.

47. Senkowski W, Gall-Mas L, Falco MM, et al. A platform for efficient establishment and drug-response profiling of high-grade serous ovarian cancer organoids. Dev Cell 2023; 58: 1106–1121 e1107.

48. Nanki Y, Chiyoda T, Hirasawa A, et al. Patient-derived ovarian cancer organoids capture the genomic profiles of primary tumours applicable for drug sensitivity and resistance testing. Sci Rep 2020; 10: 12581.

49. Maru Y, Tanaka N, Itami M, et al. Efficient use of patient-derived organoids as a preclinical model for gynecologic tumors. Gynecol Oncol 2019; 154: 189–198.

50. Hoffmann K, Berger H, Kulbe H, et al. Stable expansion of high-grade serous ovarian cancer organoids requires a low-Wnt environment. EMBO J 2020; 39: e104013.

51. Maenhoudt N, Defraye C, Boretto M, et al. Developing Organoids from Ovarian Cancer as Experimental and Preclinical Models. Stem Cell Reports 2020; 14: 717–729.

52. Thorel L, Dolivet E, Morice PM, et al. Long-term patient-derived ovarian cancer organoids closely recapitulate tumor of origin and clinical response. J Exp Clin Cancer Res 2025; 44: 282.

53. Sato S, Itamochi H, Kigawa J, et al. Combination chemotherapy of oxaliplatin and 5-fluorouracil may be an effective regimen for mucinous adenocarcinoma of the ovary: a potential treatment strategy. Cancer Sci 2009; 100: 546–551.

54. Gore M, Hackshaw A, Brady WE, et al. An international, phase III randomized trial in patients with mucinous epithelial ovarian cancer (mEOC/GOG 0241) with long-term follow-up: and experience of conducting a clinical trial in a rare gynecological tumor. Gynecol Oncol 2019; 153: 541–548.

55. Kurnit KC, Sinno AK, Fellman BM, et al. Effects of Gastrointestinal-Type Chemotherapy in Women With Ovarian Mucinous Carcinoma. Obstetrics and gynecology 2019; 134: 1253–1259.

56. Schlappe BA, Zhou QC, O’Cearbhaill R, et al. A descriptive report of outcomes of primary mucinous ovarian cancer patients receiving either an adjuvant gynecologic or gastrointestinal chemotherapy regimen. Int J Gynecol Cancer 2019.

57. Shimizu Y, Nagata H, Kikuchi Y, et al. Cytotoxic agents active against mucinous adenocarcinoma of the ovary. Oncol Rep 1998; 5: 99–101.

58. Ueda Y, Miyatake T, Nagamatsu M, et al. A phase II study of combination chemotherapy using docetaxel and irinotecan for TC-refractory or TC-resistant ovarian carcinomas (GOGO-OV2 study) and for primary clear or mucinous ovarian carcinomas (GOGO-OV3 Study). Eur J Obstet Gynecol Reprod Biol 2013; 170: 259–263.

59. Vias M, Morrill Gavarró L, Sauer CM, et al. High-grade serous ovarian carcinoma organoids as models of chromosomal instability. Elife 2023; 12.

60. Azambuja E, Durbecq V, Rosa DD, et al. HER-2 overexpression/amplification and its interaction with taxane-based therapy in breast cancer. Annals of Oncology 2008; 19: 223–232.

61. Camerini A, Donati S, Viacava P, et al. Evaluation of HER2 and p53 expression in predicting response to docetaxel-based first-line chemotherapy in advanced breast cancer. J Exp Clin Cancer Res 2011; 30: 38.

62. de Hoon JP, Veeck J, Vriens BE, et al. Taxane resistance in breast cancer: a closed HER2 circuit? Biochim Biophys Acta 2012; 1825: 197–206.

63. Meagher NS, Gorringe KL, Wakefield M, et al. Gene-Expression Profiling of Mucinous Ovarian Tumors and Comparison with Upper and Lower Gastrointestinal Tumors Identifies Markers Associated with Adverse Outcomes. Clin Cancer Res 2022; 28: 5383–5395.

