## Supplementary Information for "Comprehensive drug efficacy data for mucinous ovarian carcinoma using a novel and extensive biobank of patient-derived organoid models"

**Supplementary Figures:**

**Supplementary Figure 1 Media optimisation.** The **Table** shows the constituents of the various media tested. CM - conditioned media for drug screening and other experiments in the study, however, recombinant protein used in initial testing. *Y27632 could be removed once the line was established but was always kept in for the first several passages. **A.** CRC medium compared with addition of insulin, B-estradiol and progesterone in ORG03 (2 wells per condition). None of the additives improved the survival of organoids over time. **B**. Comparison of CRC with ORGM1 in ORG09 showing marginal improvement in organoid number with the additon of recombinant RSPO1 and Wnt-3a. **C**. Comparison of ORGM0 and ORGM1 with various additives, with the addition of Noggin substantially increasing organoid number. **D**. MDS plot and volcano plot of organoid RNAseq comparing those grown in Kopper (Hans) with ORG3M (ORG). Only a single gene was statistically significant. MDS plot corrected for patient ID.

**Supplementary Figure 2 Media testing.** Six organoid lines were grown in media lacking Wnt3a, Noggin, FGF10, RSPO1, none of Wnt/Noggin/RSPO or with reduced Forskolin (50%). 3-6 wells for each condition in a 384-well plate were imaged at the timepoint indicated (in days) and objects were counted in Qupath. The average number relative to complete media is shown with standard error of the mean. Dots are counts for each well. Scale bar 200 pixels, approximately 12.5% the diameter of the well at the point of imaging.

**Supplementary Figure 3 Organoid cryopreservation**. **A** Organoids were cryopreserved in different cryopreservation media with Rho kinase inhibitors before being re-established after three months and imaged on day 10. **B** Representative brightfield images of ORG38 after revival from cryopreservation when in exponential phase or passage endpoint in different freezing media (protocols 1-5). Scale bar: 2000 µm. **C-D** The average number of organoids per well after cryopreservation in exponential phase (C) and passage endpoint (D). Data are mean ± SD with N = 3 wells per condition

**Supplementary Figure 4 Individual organoid morphologies.** Images taken on the Cytation 5 or 10 Multimode Imager at 4X magnification, except for ORG16 and ORG41 taken on the Evos Fl Microscope at 4X magnification

**Supplementary Figure 5 Organoid immunohistochemistry of matched organoids and parental tumours.** **IHC concordance** between organoids and tumours by A) case and B) marker. Discordance is when the staining cannot be easily explained by heterogeneity e.g. positive vs negative, whereas partial concordance could be explained by heterogeneity e.g. focal positive vs patchy positive. Following pages show individual cases. Stained slides were scanned at 40X magnification on the VS120 Slide Scanner by Olympia and screenshots taken of representative fields.

**Supplementary Figure 6 DNA sequencing circos and copy number plots**. Outputs from Oncoanalyser showing two circos plots for each sample analysed by whole genome sequencing. At left is the ploidy corrected copy number with structural variants. Green – copy number gains, red – copy number losses, blue – allelic imbalance, orange – loss of heterozygosity, dots – variant allele frequency. At right is the uncorrected copy number (blue) and B-allele frequency (orange) plots. For Org16, output is from CopywriteR, for ORG44 output from PureCN and ORG76 output from CNVkit are shown.

**Supplementary Figure 7 Drug screening individual dose response curves. A**. Dose curves of organoid lines treated with two doses. Shown is the average CTG values of 2 wells, normalised to control wells, from each of duplicate experiments with the standard error of the mean, except for ORG46, ORG49, ORG60 and ORG73 which are only a single biological replicate (2 wells). **B**. As for A except treated with a single dose.

**Supplementary Figure 8 Comparison between 1 and 2 dose experiments**. Relative AUC of all cases, comparing 1 dose and 2 doses. Dose response curves for ORG49 and ORG73 for all chemotherapy drugs, which each had one experiment with a single dose and one with two doses. Error bars represent the range of two wells.

**Supplementary Figure 9 Synergy drug screening**. Cell viability as measured by CTG relative to control wells. Data points the average of two biological replicates, each the average of two wells. Error bars are the range.

**Supplementary Figure 10. Early passage screening for clinical feasibility.** Dose response curves of CTG values normalised to control wells using a refined set of drugs compared to the previous screen (docetaxel and cisplatin removed, SN38 added). Note that additional lower doses were added for paclitaxel and gemcitabine, while higher doses were added for carboplatin and oxaliplatin. Average and range of two wells. **A**. ORG77. **B**. ORG78

**Supplementary Tables.**

**Supplementary Table 1 Clinicopathological information from all organoid cases**

**Supplementary Table 2 Immunohistochemistry scoring for organoid cases**

**Supplementary Table 3 DNA and RNA sequencing metrics**

**Supplementary Table 4 STR profiles**

**Supplementary Table 5 High confidence variant information**

**Supplementary Table 6 Copy number segments**

**Supplementary Table 7 Differentially expressed genes**

**Supplementary Table 8a Drug response data**

**Supplementary Table 8b Drug AUC values**

**Supplementary Table 9 Correlations between drug responses and tumour/organoid features**

**Supplementary Methods**

1. **Media formulation**

| **Components** | **Kopper** | **ORG-M3** | **Wash media** | **Company** | **Catalog number** |
| --- | --- | --- | --- | --- | --- |
| Base media | DMEM/F12 | DMEM/F12 | DMEM/F12 | Sigma-Aldrich | D6421 |
| Pen/Strep | 50U/mL; 50ug/mL | 50U/mL; 50ug/mL | 50U/mL; 50ug/mL | Gibco | 15070063 |
| Hepes | 2 mM | 2 mM | 10 mM | Gibco | 15630080 |
| Glutamax | 1x | 1x | 1x | Gibco | 35050061 |
| Nicotinamide | 10mM | 10mM |  | Sigma-Aldrich | N06036 |
| A8301 | 0.5 µM | 0.5 µM |  | TOCRIS | 2939 |
| FGF 10 | 10 ng/ml | 10 ng/ml |  | Peprotech | 100-26 |
| Heregulin-β1 | 37.5 ng/ml |  |  | Peprotech | 100-03 |
| EGF | 5 ng/mL | 50 ng/mL |  | Sigma Aldrich | SRP3027 |
| Y27632 (Y27) | 5 µM | 10 µM |  | TOCRIS | RDS125450 |
| N acetyl Cysteine (NAC) | 1 mM | 1 mM |  | Sigma-Aldrich | A9165 |
| B27 | 1x | 2x |  | ThermoFisher Sc. | 17504-044 |
| R-Spondin 1 | 10% | 10% |  | in house conditioned media |  |
| NOGGIN | 10% | 10% |  | in house conditioned media |  |
| Wnt-3a | 20% | 20% |  | in house conditioned media |  |
| Β-Estradiol | 0.1 µM |  |  | Sigma-Aldrich | E8875-16 |
| Forskolin | 10 µM |  |  | TOCRIS | RDS109910 |
| Hydrocortisone | 0.5 µg/mL |  |  | Sigma-Aldrich | H0888 |
| FGF2 |  | 10 ng/ml |  | Peprotech | 100-18B |
| Gastrin |  | 1 µg/mL |  | Peprotech | 100-18B |
| N2 |  | 1x |  | Gibco | 17502048 |
