## Supplementary Figure for "Comprehensive drug efficacy data for mucinous ovarian carcinoma using a novel and extensive biobank of patient-derived organoid models"

| Component | CRC-OBM | MOC-ORGM0 | MOC-ORGM1 | MOC-ORGM2 | MOC-ORGM3 | Kopper |
|---|---|---|---|---|---|---|
| DMEM/F12 | | | | | | |
| Hepes | | | | | | |
| Glutamax | | | | | | |
| Pen-Strep | | | | | | |
| A8301 | | | | | | |
| B27 | | | | | | |
| EGF | | | | 50 ng/mL | 5 ng/mL | 5 ng/mL |
| Gastrin | | | | | | |
| NAC | | | | | | |
| Y27632 | | | | | 10 µM | 5 µM* |
| Sb202190 | | | | | | |
| Sb431542 | | | | | | |
| RSPO1 | | | | | | CM |
| Wnt-3a | | | | | | CM |
| FGF 10 | | | | | | |
| Nicotidamine | | | | | | |
| FGF 2 | | | | | | |
| Prostaglandin | | | | | | |
| Noggin | | | | | | CM |
| N2 | | | | | | |
| B-Oestradiol | | | | | | |
| Progesterone | | | | | | |
| Insulin | | | | | | |
| Heregulin-B1 | | | | | | |
| Forskolin | | | | | | |
| Hydrocortisone | | | | | | |

A B C

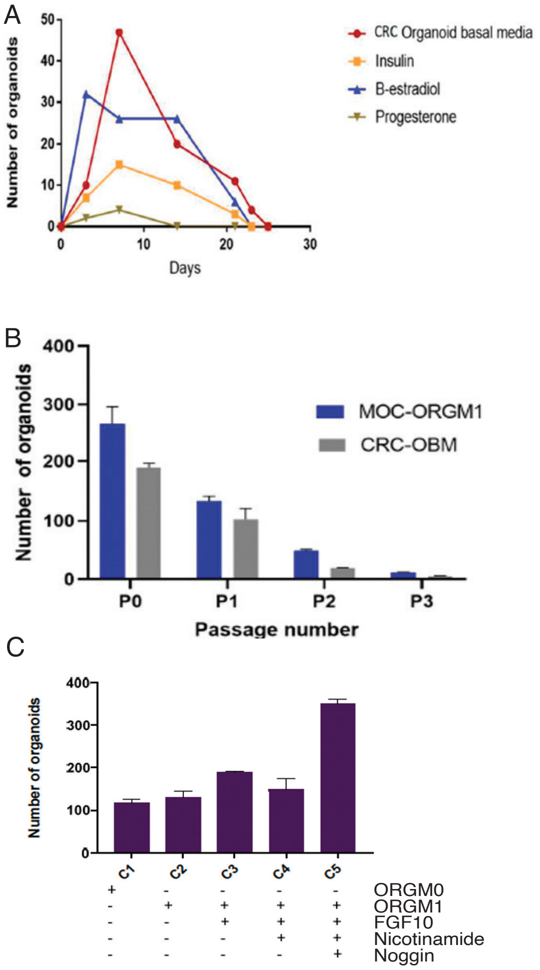


D

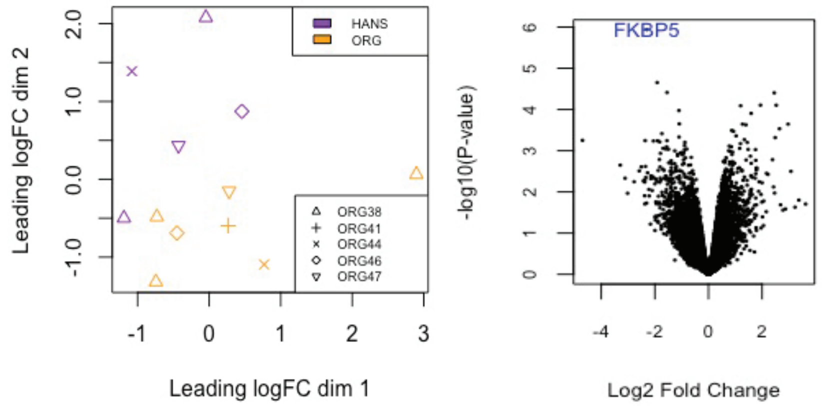


**Supplementary Figure 1.** The **Table** shows the consituents of the various media tested. CM - conditioned media for drug screening and other experiments in the study, however, recombinant protein used in initial testing. *Y27632 could be removed once the line was established but was always kept in for the first several passages. **A.** CRC medium compared with addition of insulin, B-estradiol and progesterone in ORG03 (2 wells per condition). None of the additives improved the survival of organoids over time. **B**. Comparison of CRC with ORGM1 in ORG09 showing marginal improvement in organoid number with the additon of recombinant RSPO1 and Wnt-3a. **C**. Comparison of ORGM0 and ORGM1 with various additives, with the addition of Noggin substantially increasing organoid number. **D**. MDS plot and volcano plot of organoid RNAseq comparing those grown in Kopper (Hans) with ORG3M (ORG). Only a single gene was statistically significant. MDS plot corrected for patient ID.

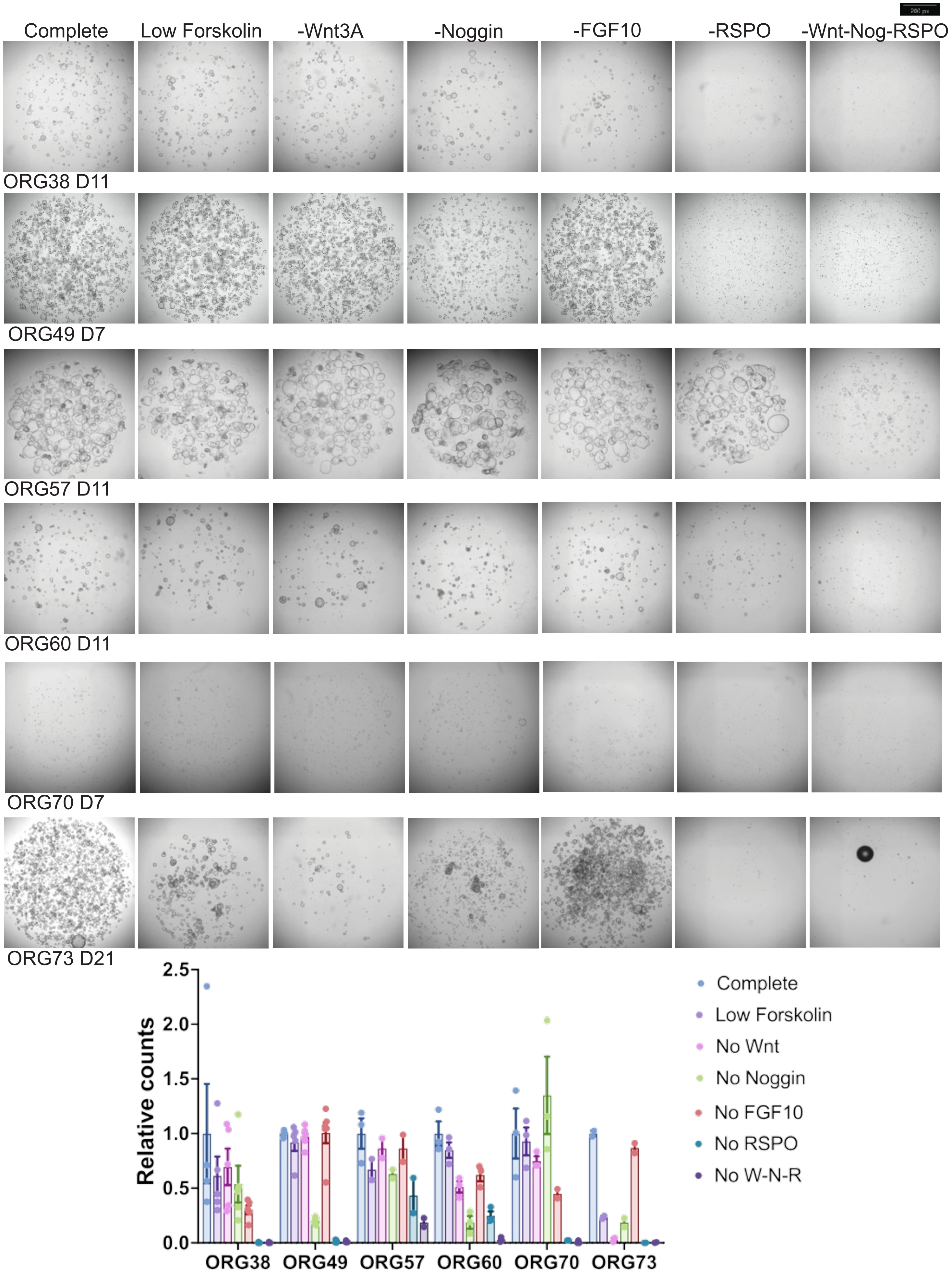


**Supplementary Figure 2. Media testing.** Six organoid lines were grown in media lacking Wnt3a, Noggin, FGF10, RSPO1, none of Wnt/Noggin/RSPO or with reduced Forskolin (50%). 3-6 wells for each condition in a 384-well plate were imaged at the timepoint indicated (in days) and objects were counted in Qupath. The average number relative to complete media is shown with standard error of the mean. Dots are counts for each well. Scale bar 200 pixels, approximately 12.5% the diameter of the well at the point of imaging.

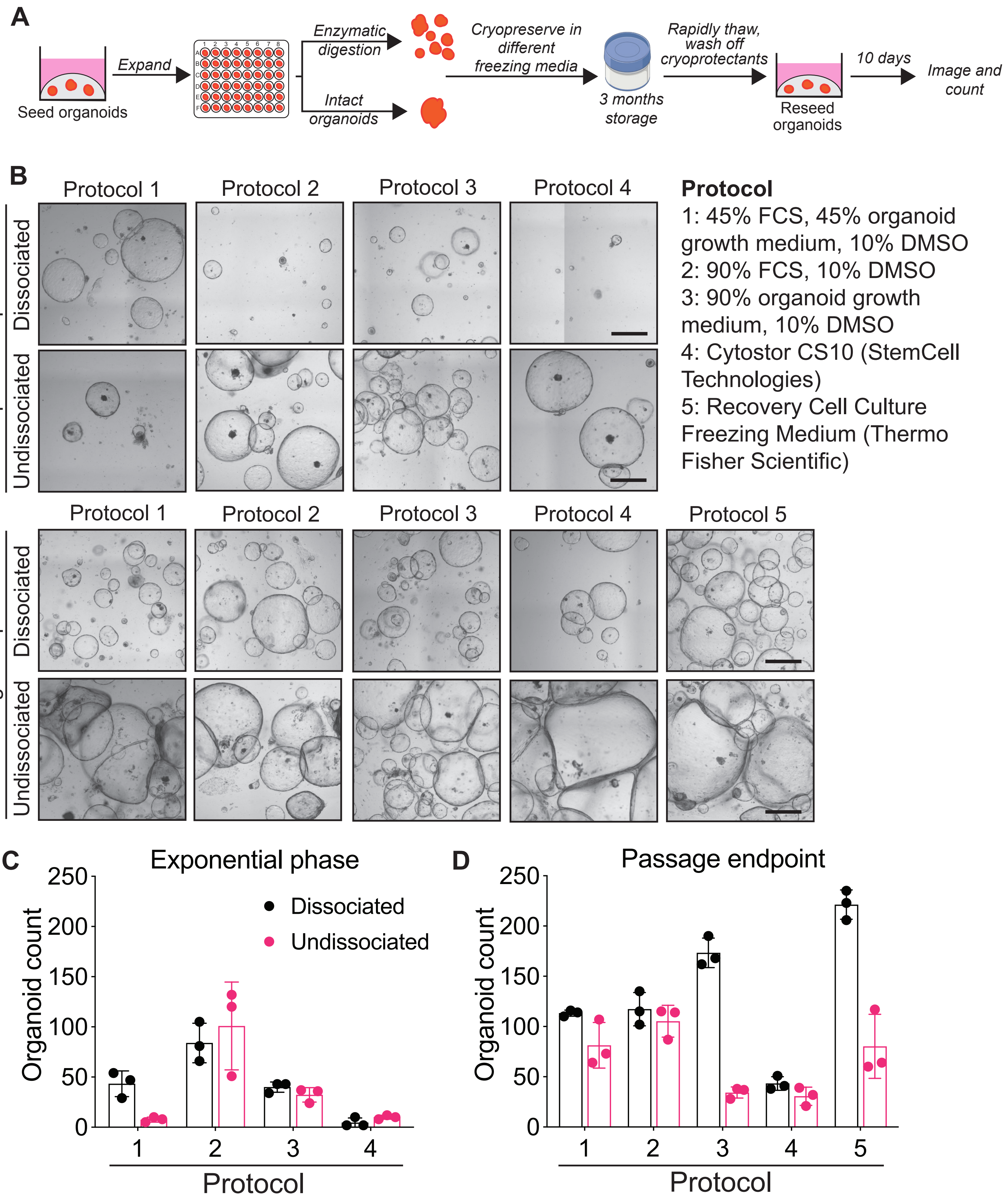


**Supplementary Figure 3. A** Organoids were cryopreserved in different cryopreservation media with Rho kinase inhibitors before being re-established after three months and imaged on day 10. **B** Representative brightfield images of ORG38 after revival from cryopreservation when in exponential phase or passage endpoint in different freezing media (protocols 1-5). Scale bar: 2000 µm. **C-D** The average number of organoids per well after cryopreservation in exponential phase (C) and passage endpoint (D). Data are mean ± SD with N = 3 wells per condition

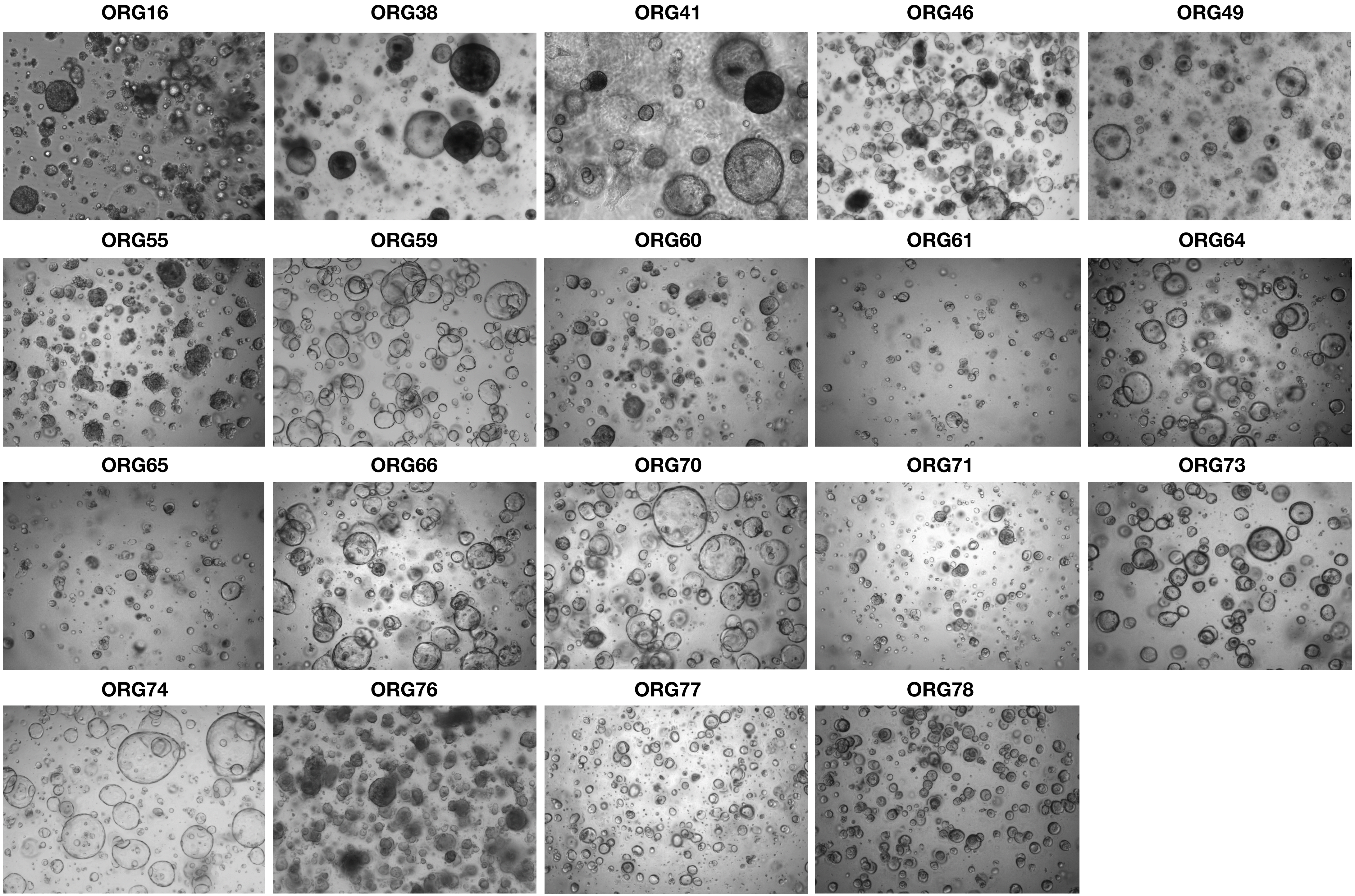


**Supplementary Figure 4. Examples of varying organoid morphologies for each line**

Images taken on the Cytation 5 or 10 Multimode Imager at 4X magnification, except for ORG16 and ORG41 taken on the Evos Fl Microscope at 4X magnification

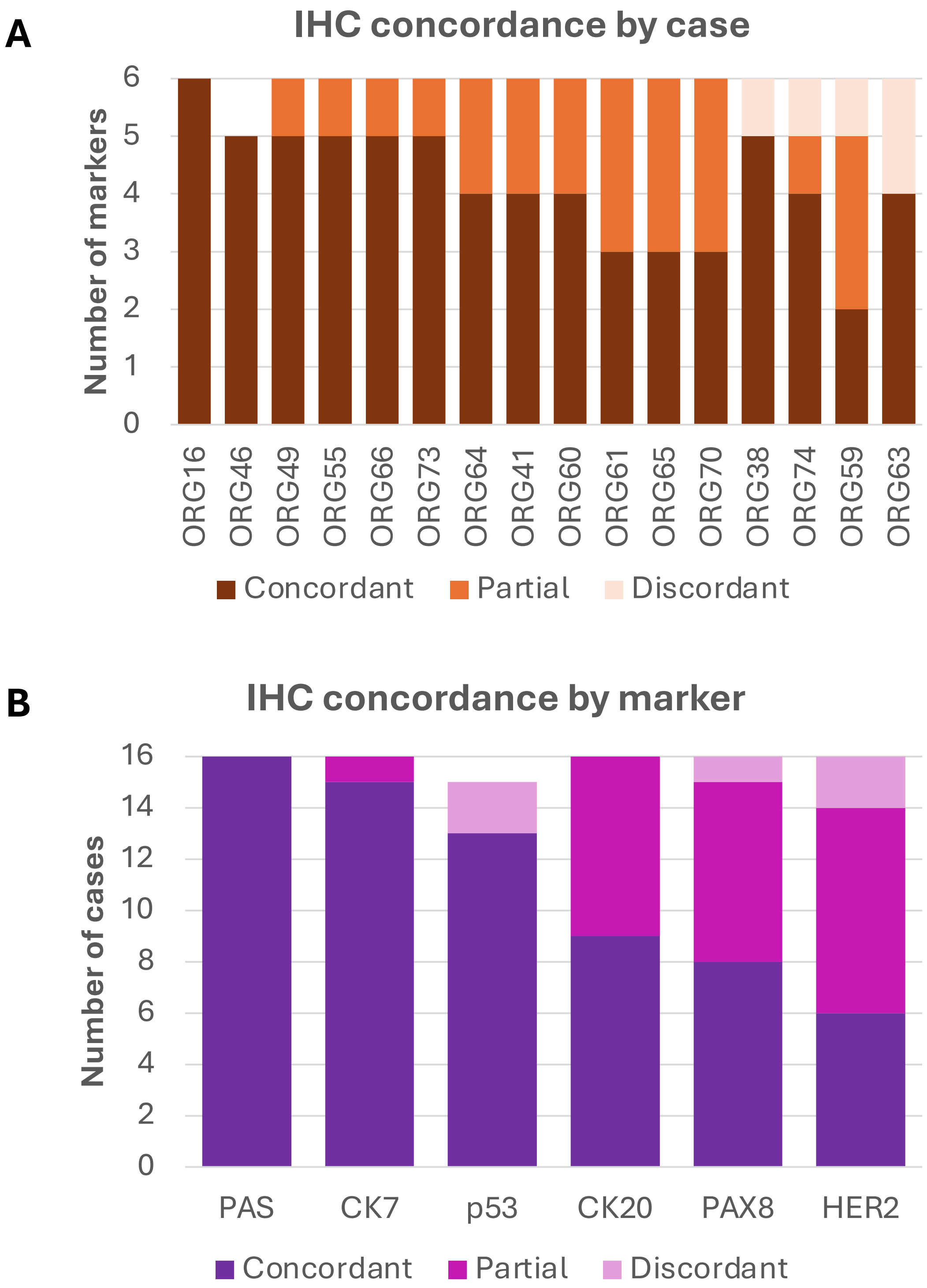


**Supplementary Figure 5 Organoid immunohistochemistry of matched organoids and parental tumours. IHC concordance** between organoids and tumours by A) case and B) marker. Discordance is when the staining cannot be easily explained by heterogeneity e.g. positive vs negative, whereas partial concordance could be explained by heterogeneity e.g. focal positive vs patchy positive. Following pages show individual cases. Stained slides were scanned at 40X magnification on the VS120 Slide Scanner by Olympia and screenshots taken of representative fields.

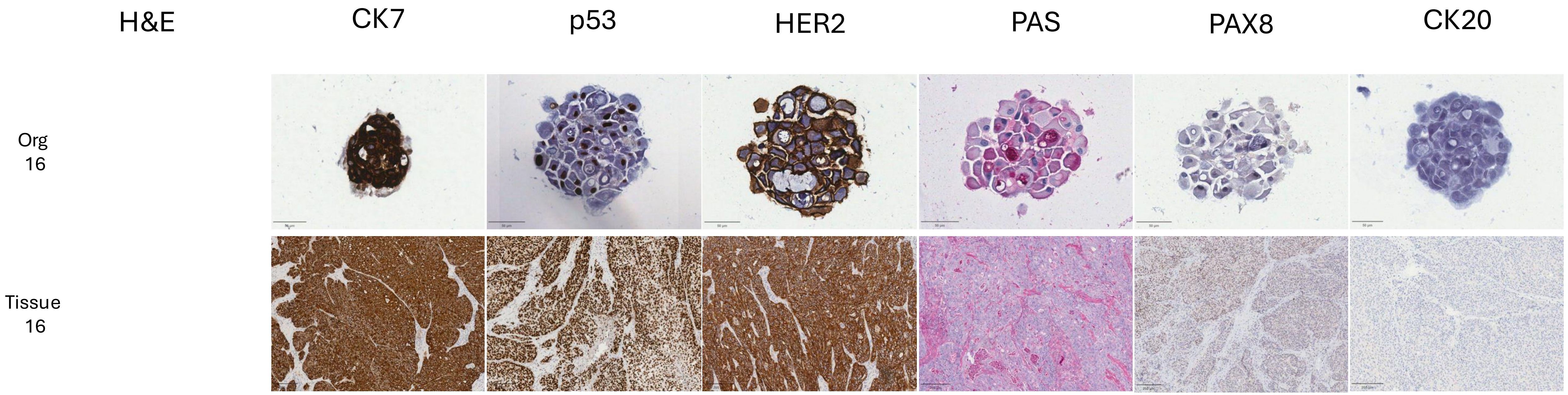
H&E
CK7
p53
HER2
PAS
PAX8
CK20
Org
16
Tissue
16

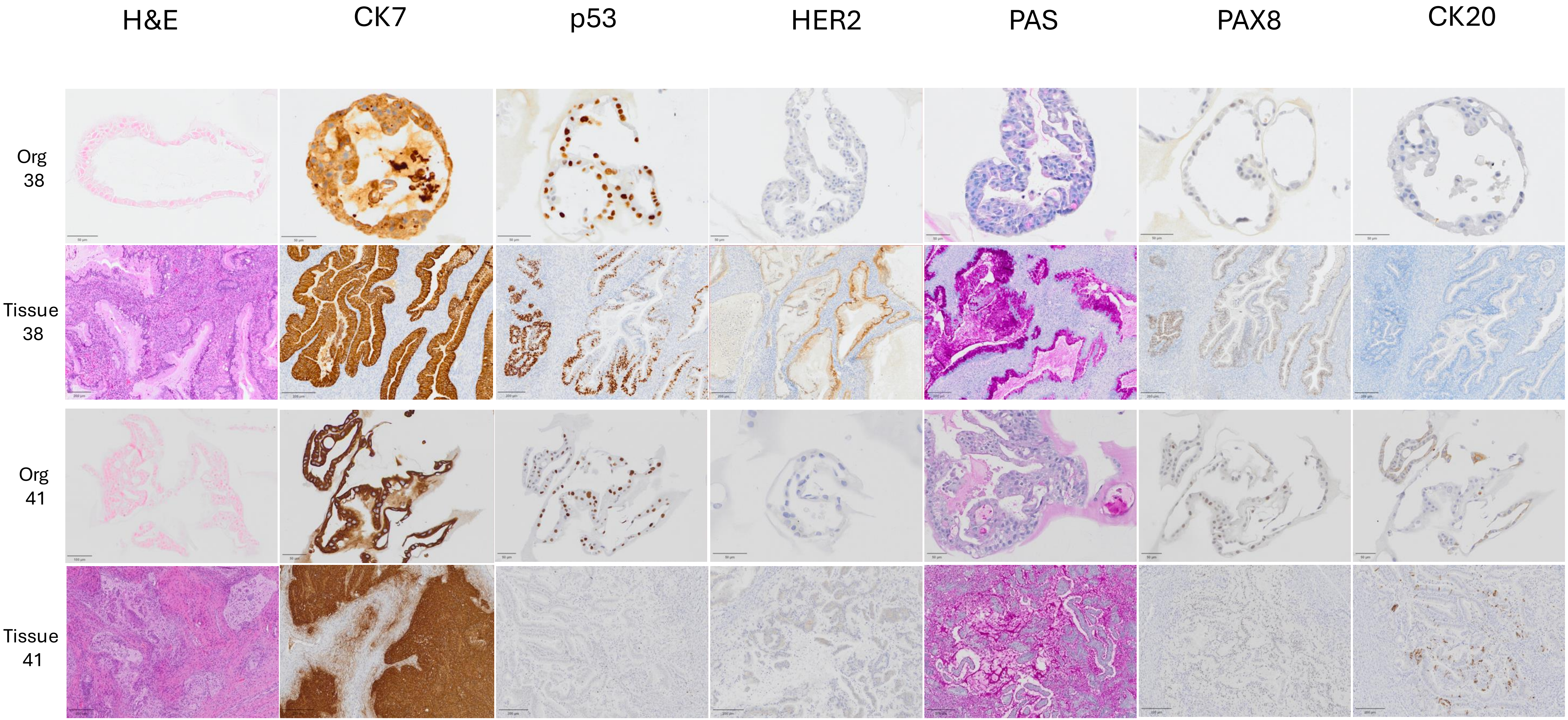
H&E
CK7
p53
HER2
PAS
PAX8
CK20
Org 38
Tissue 38
Org 41
Tissue 41

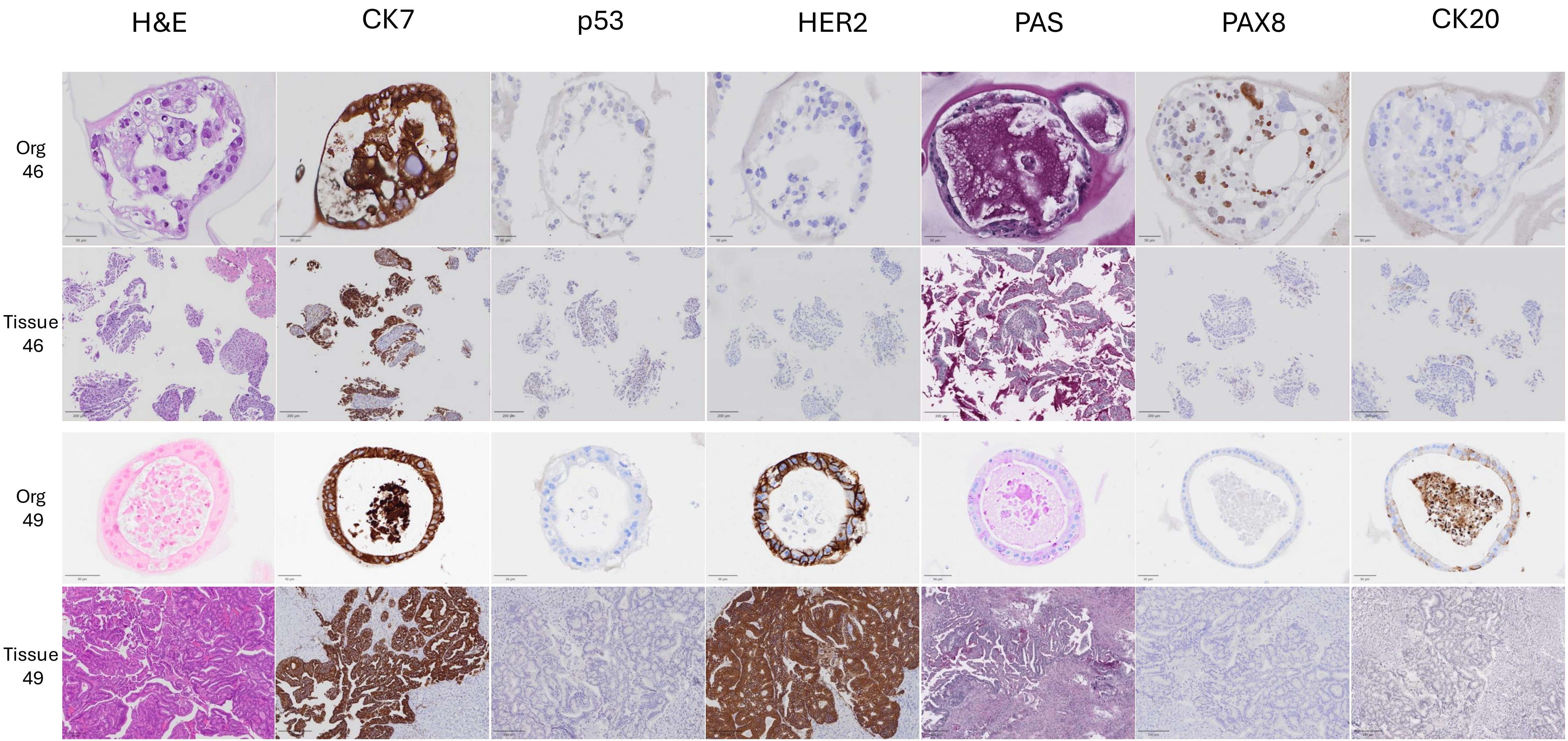
H&E
CK7
p53
HER2
PAS
PAX8
CK20
Org 46
Tissue 46
Org 49
Tissue 49

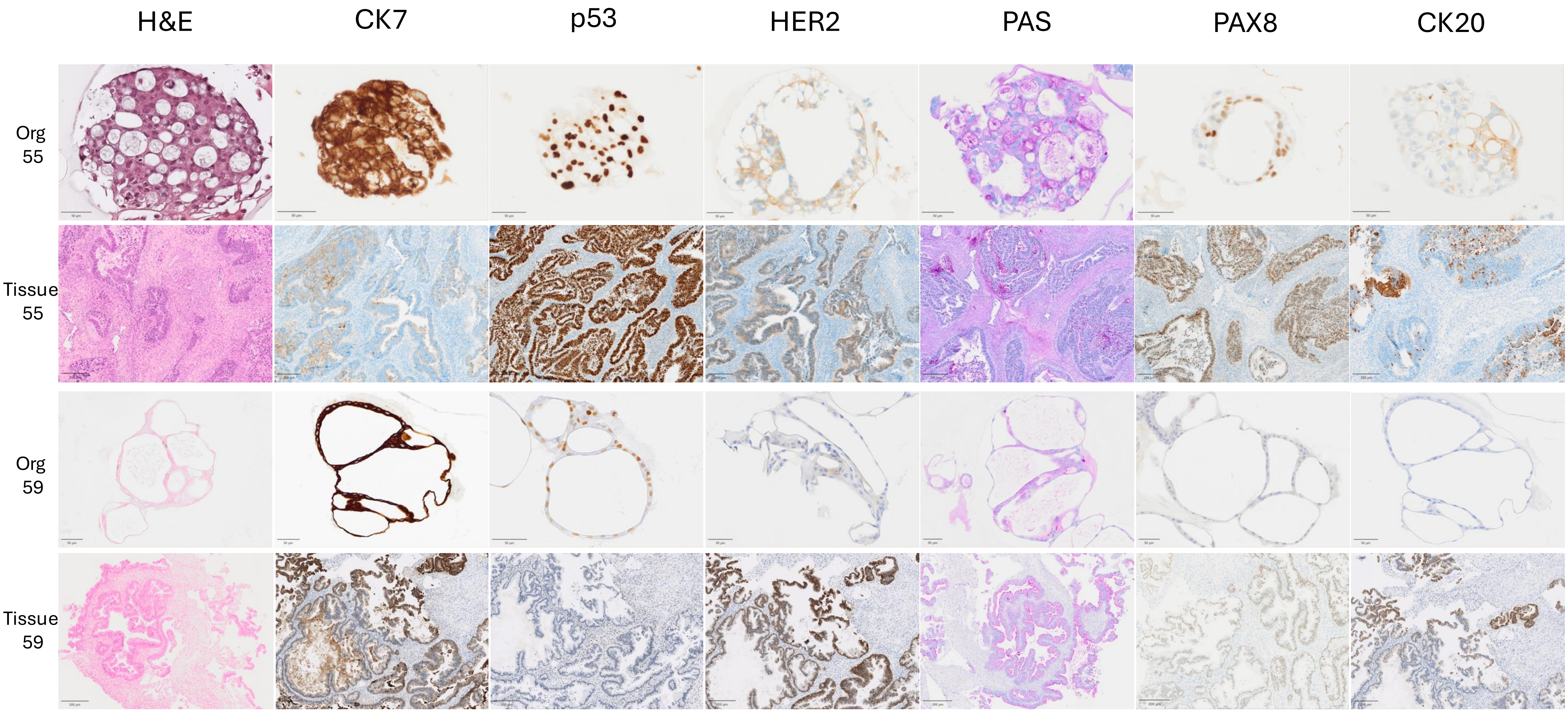
H&E
CK7
p53
HER2
PAS
PAX8
CK20
Org 55
Tissue 55
Org 59
Tissue 59

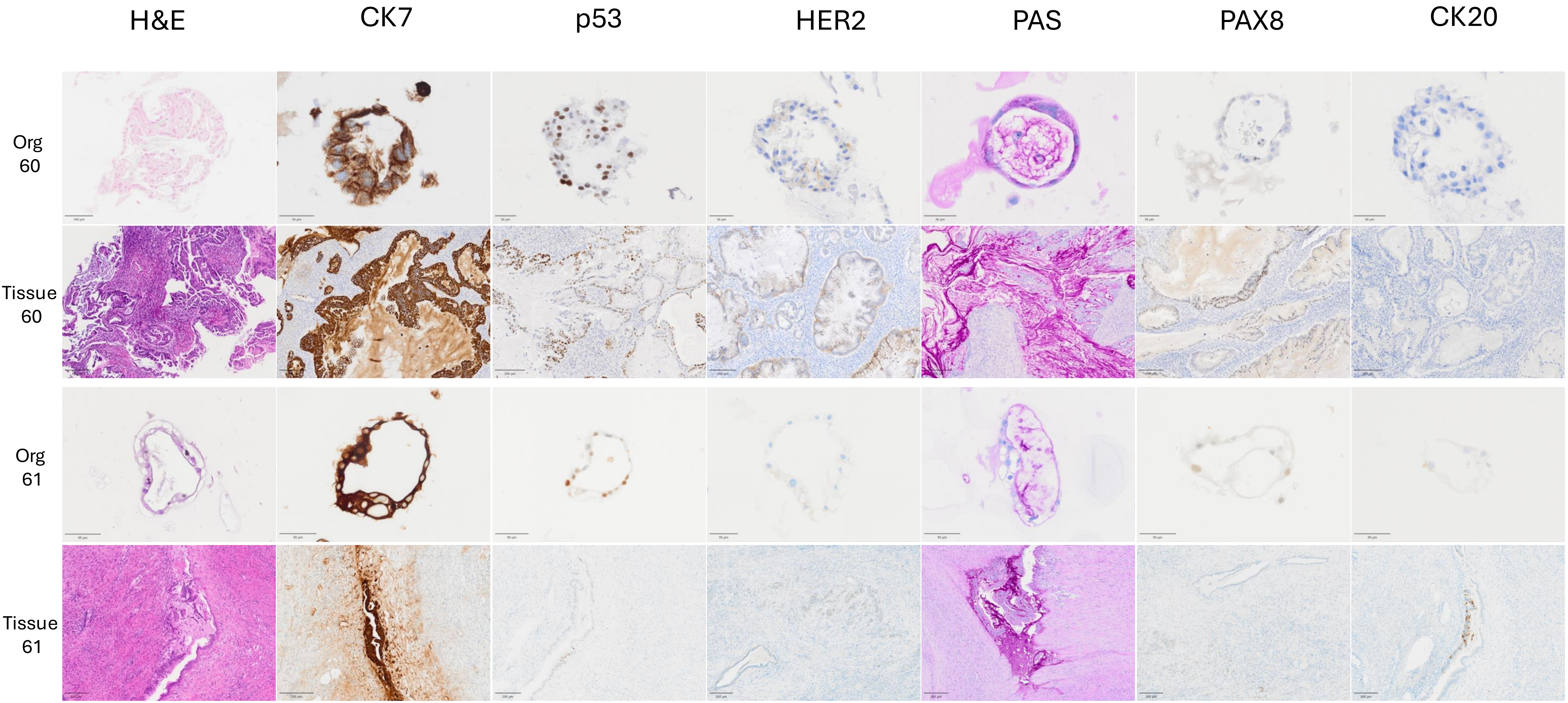
H&E
CK7
p53
HER2
PAS
PAX8
CK20
Org 60
Tissue 60
Org 61
Tissue 61

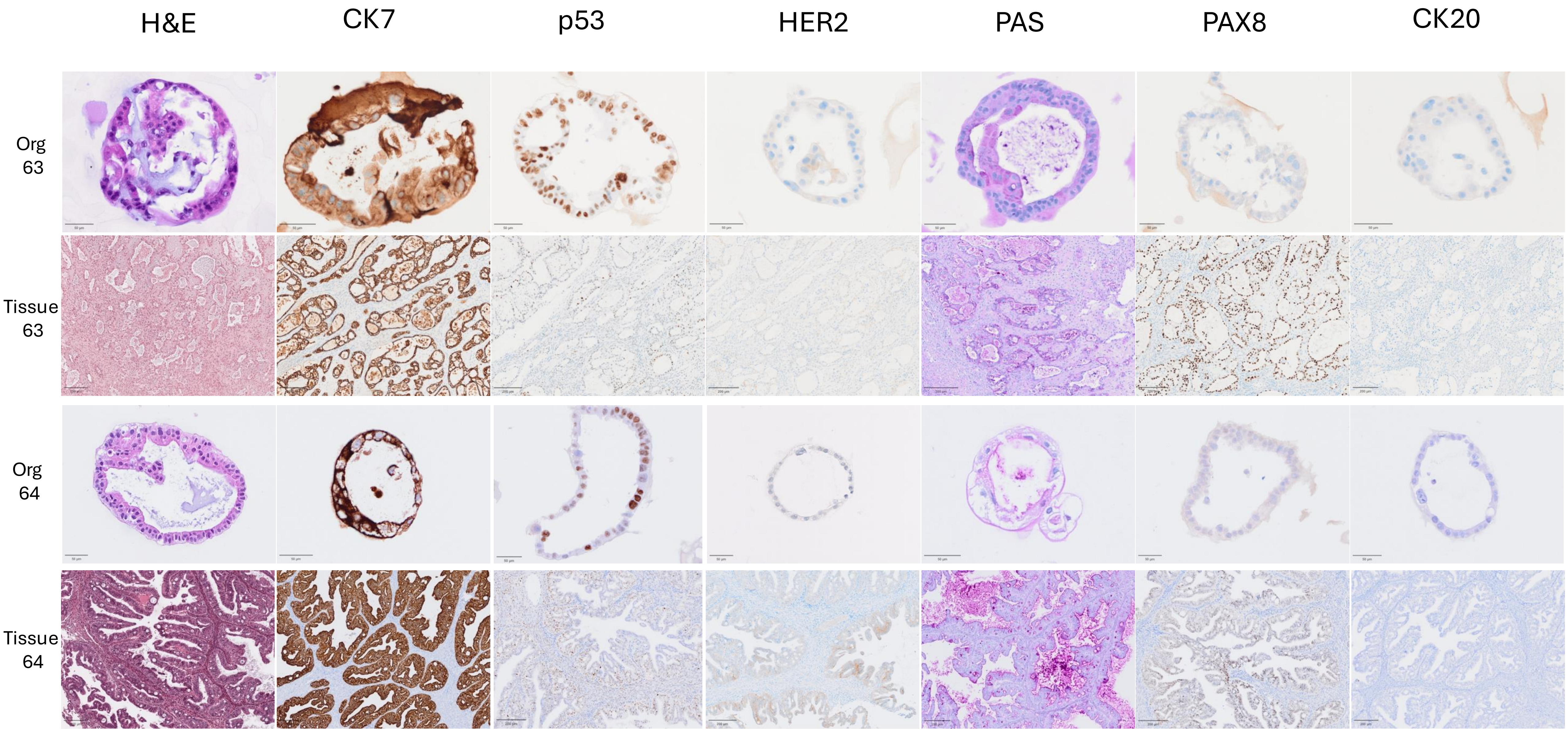
H&E
CK7
p53
HER2
PAS
PAX8
CK20
Org 63
Tissue 63
Org 64
Tissue 64

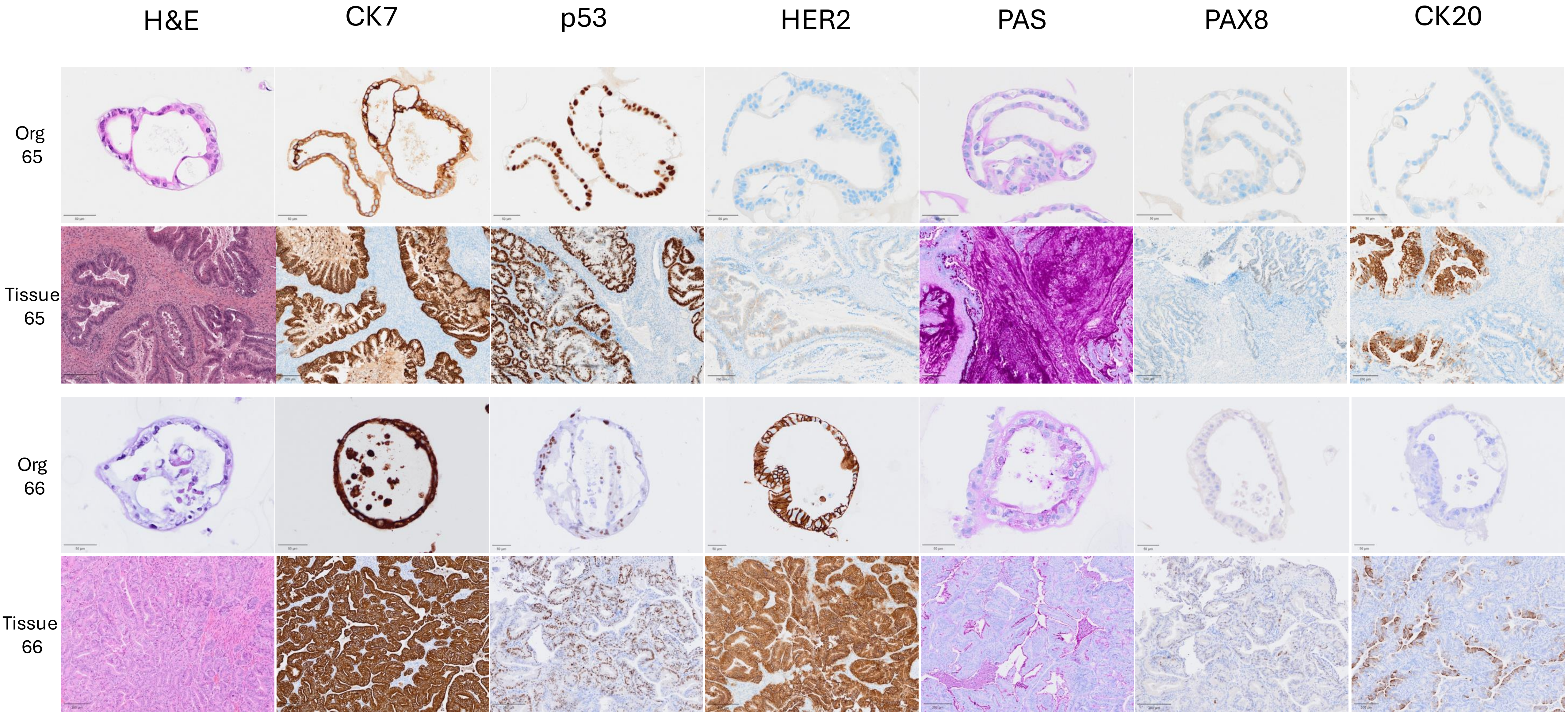
H&E
CK7
p53
HER2
PAS
PAX8
CK20
Org 65
Tissue 65
Org 66
Tissue 66

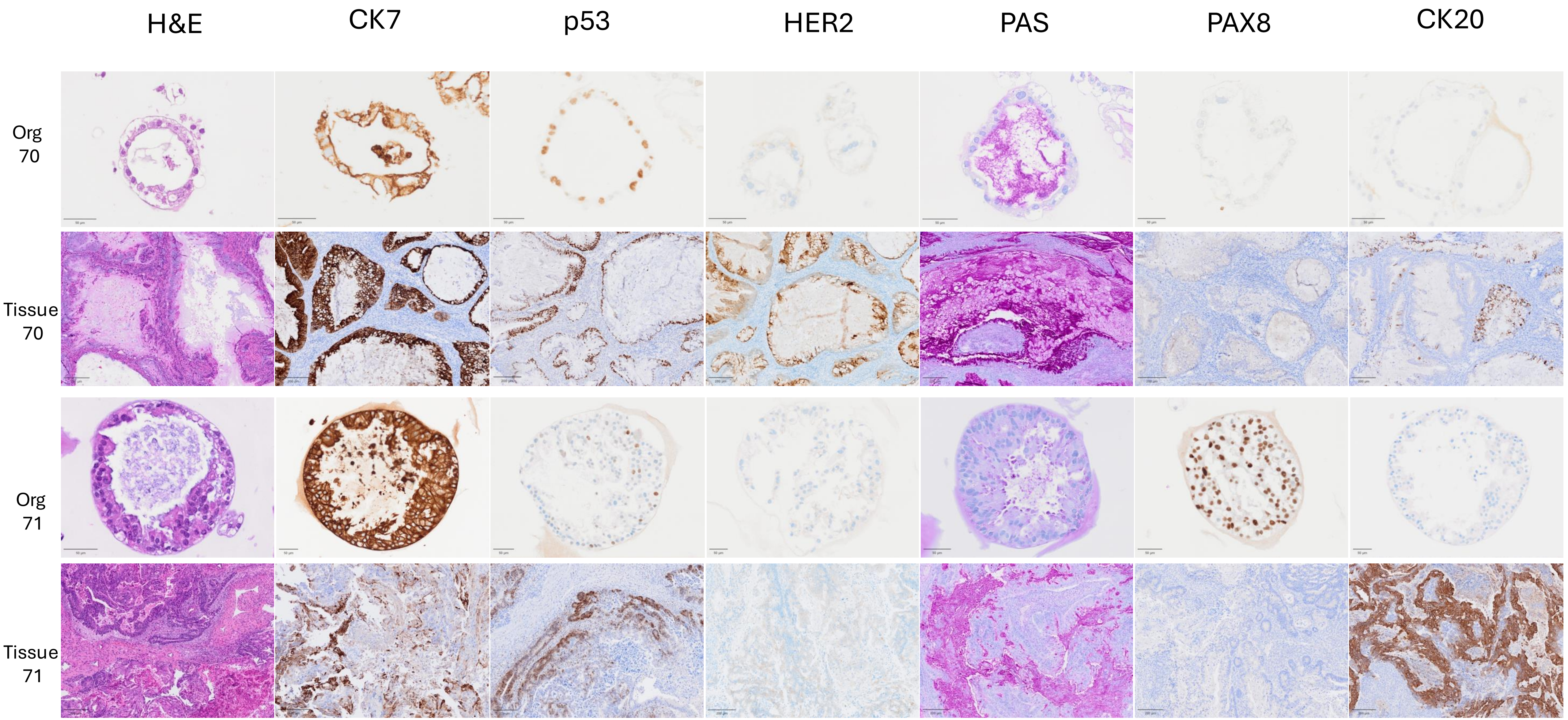
H&E
CK7
p53
HER2
PAS
PAX8
CK20
Org 70
Tissue 70
Org 71
Tissue 71

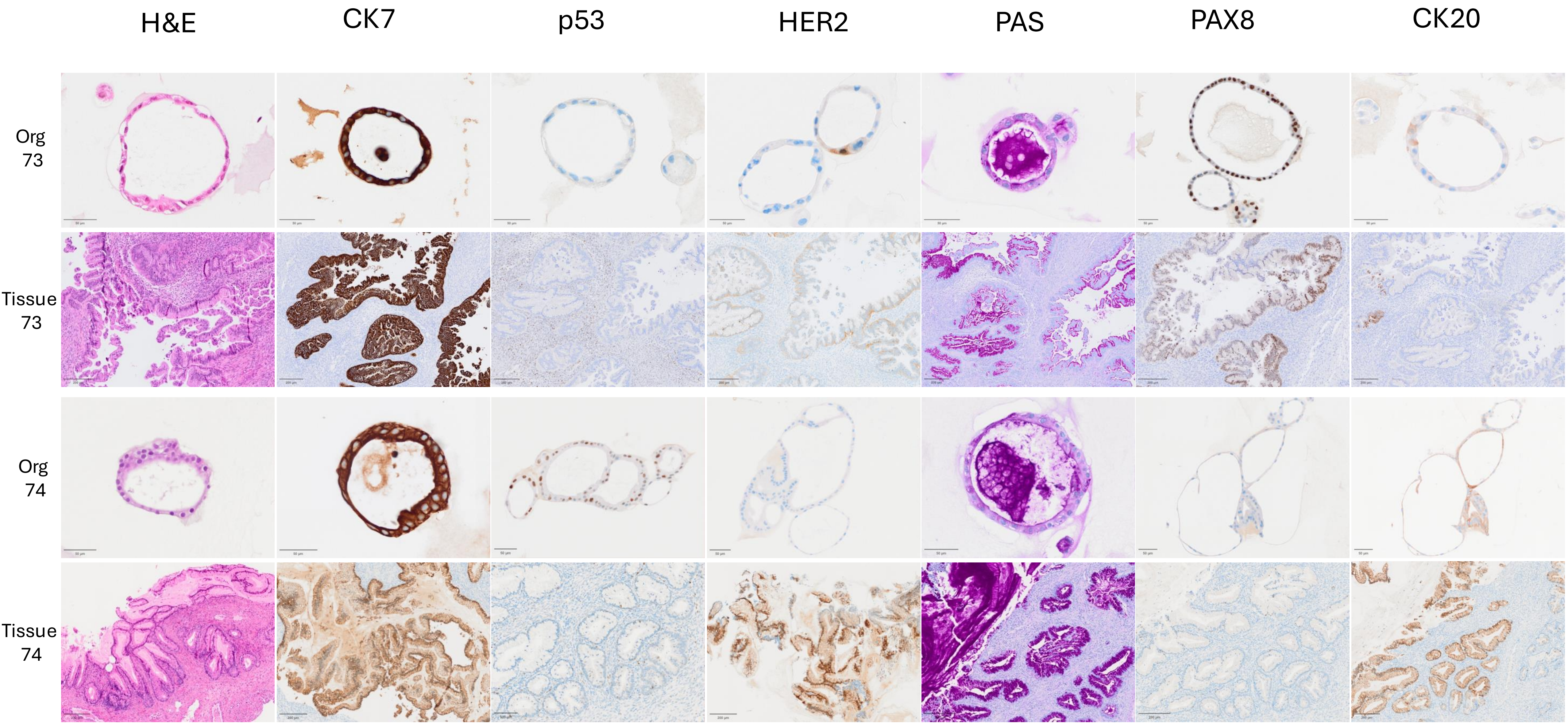
H&E
CK7
p53
HER2
PAS
PAX8
CK20
Org 73
Tissue 73
Org 74
Tissue 74

H&E CK7 p53 HER2 PAS PAX8 CK20

Org 76

Tissue 76

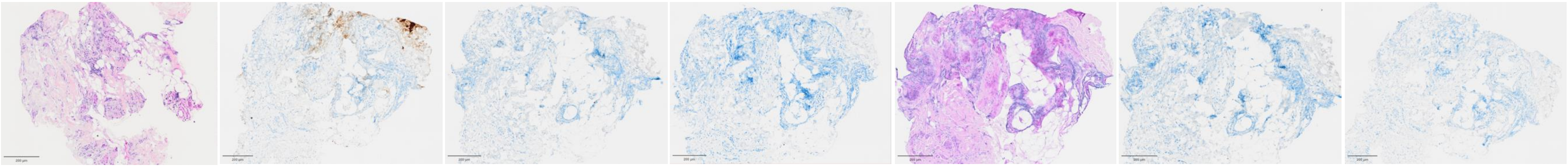

ORG38T

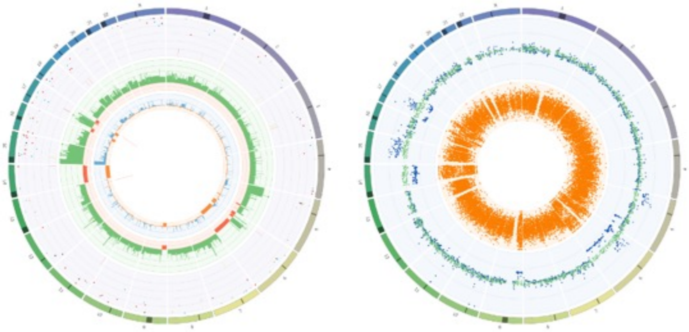

ORG38Org

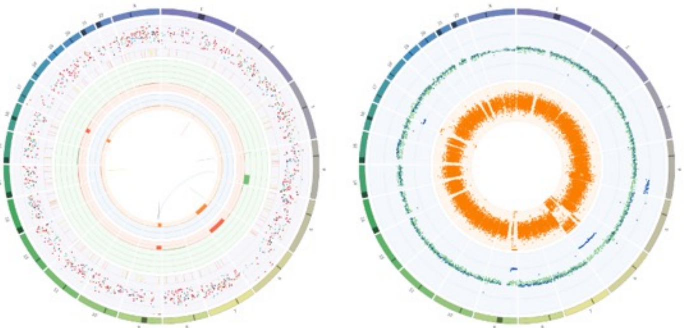

ORG41T

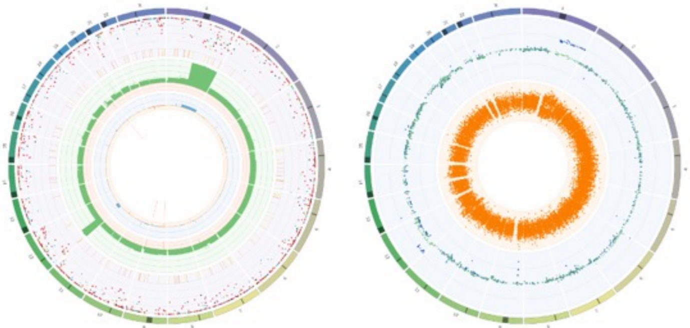

ORG41Org

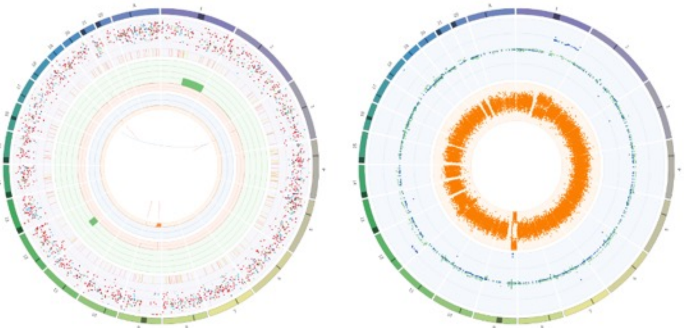

ORG46T

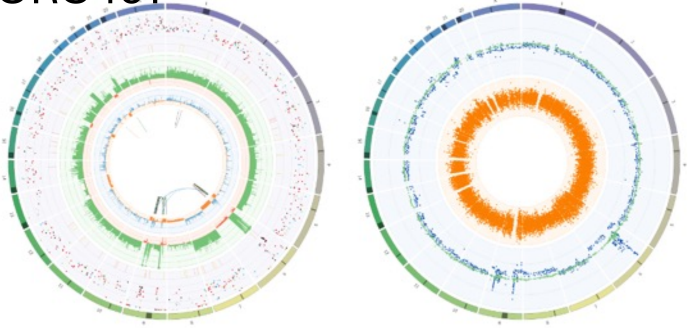

ORG46Org

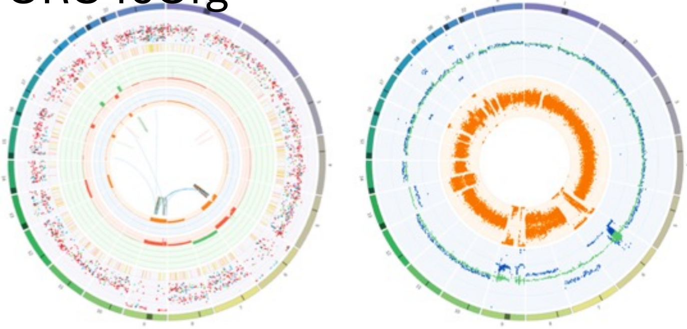

ORG49T

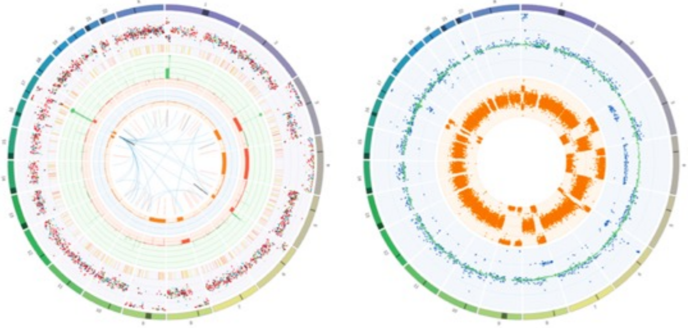

ORG49Org

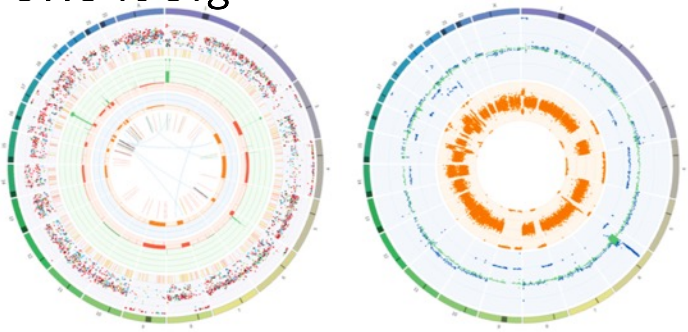

ORG55T

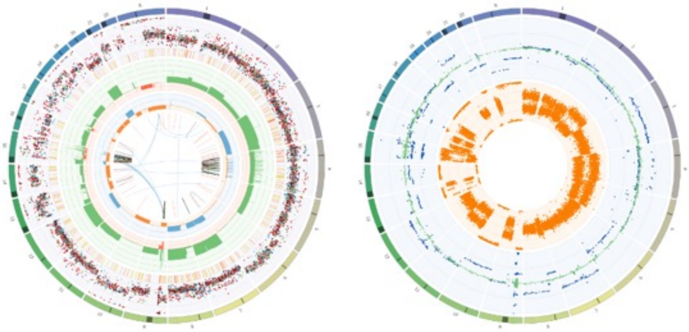

ORG55Org

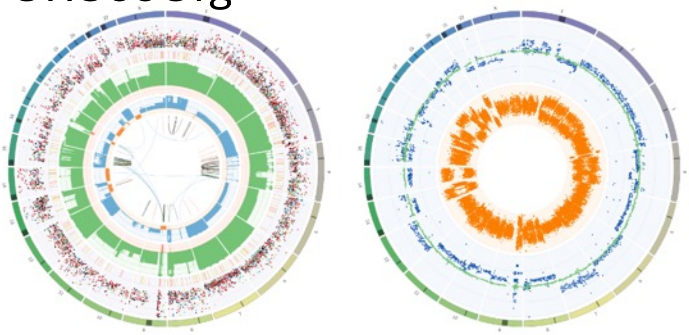

ORG59T (unmatched)

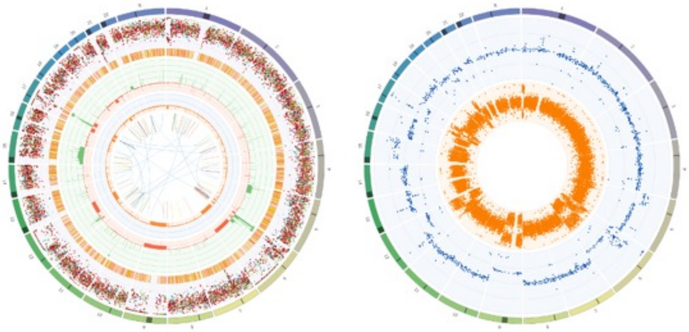

ORG57Org (unmatched)

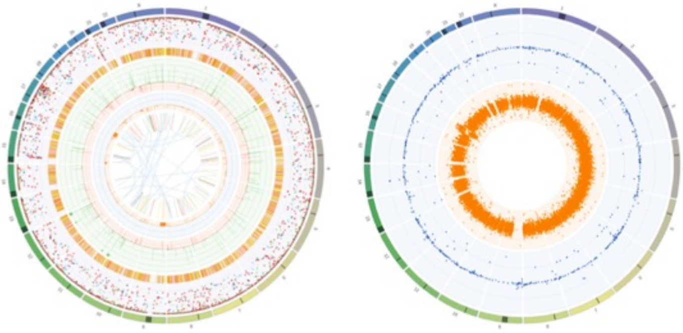

ORG60T

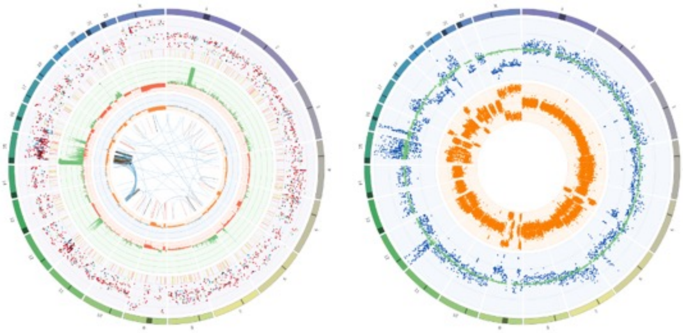

ORG60Org

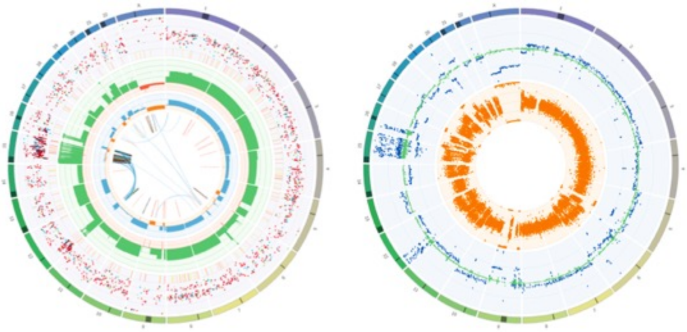

ORG63T

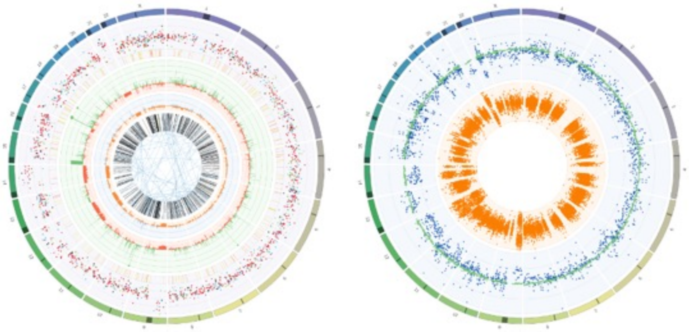

ORG63Org

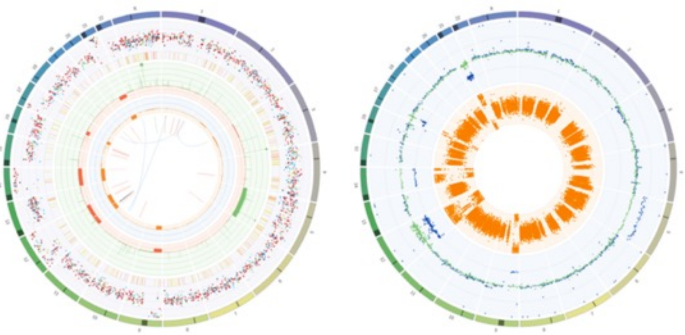

ORG64T

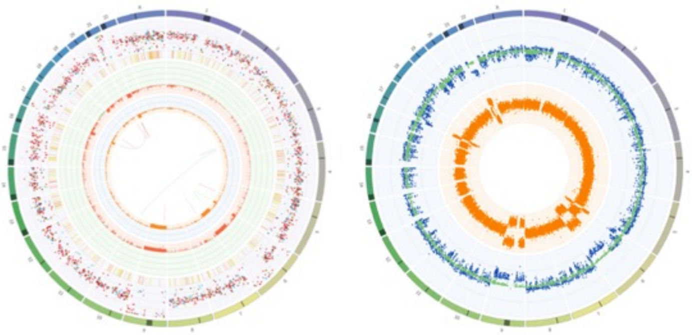

ORG64Org

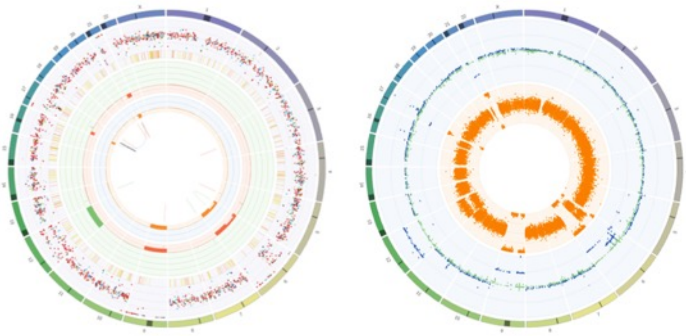

ORG65T

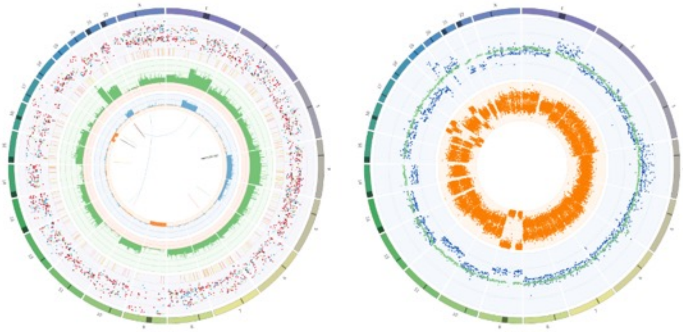

ORG65Org

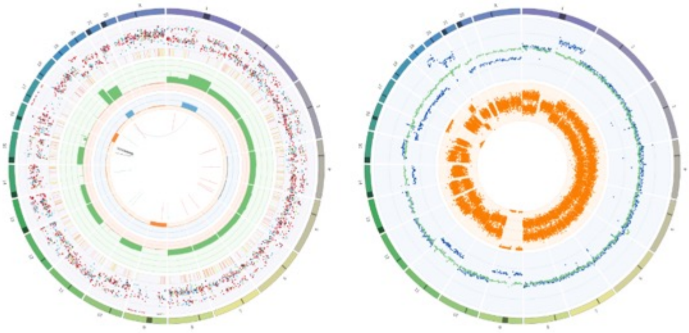

ORG61Org (unmatched)

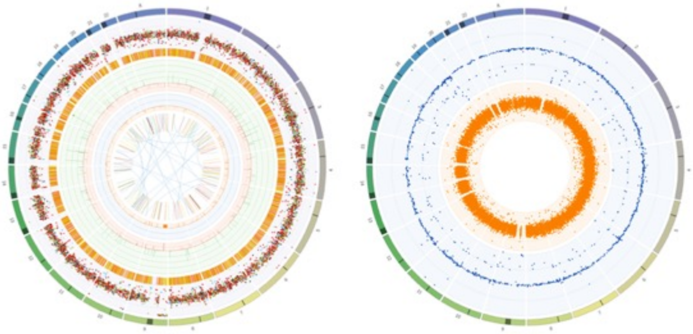

ORG62Org (unmatched)

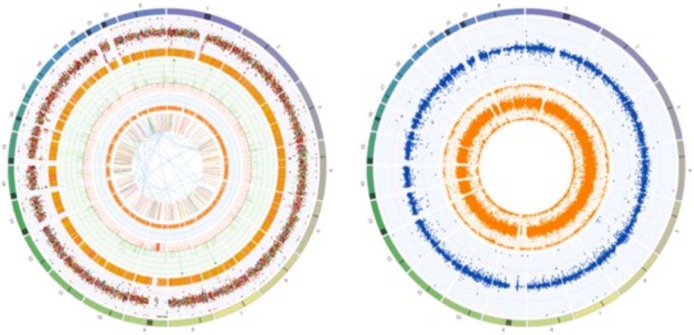

ORG66T

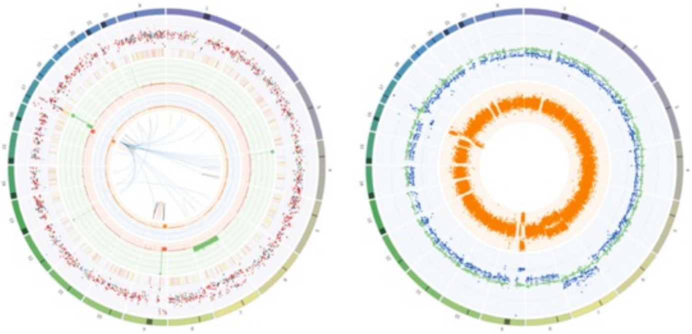

ORG66Org

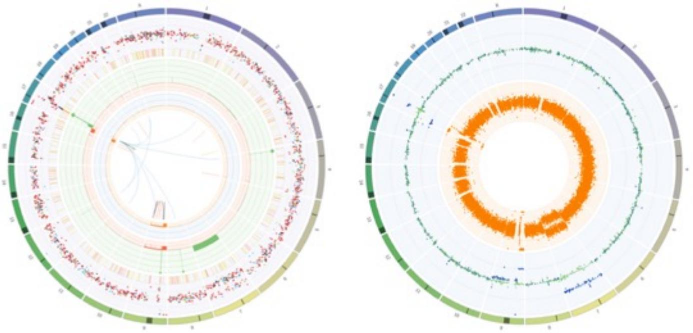

ORG70T

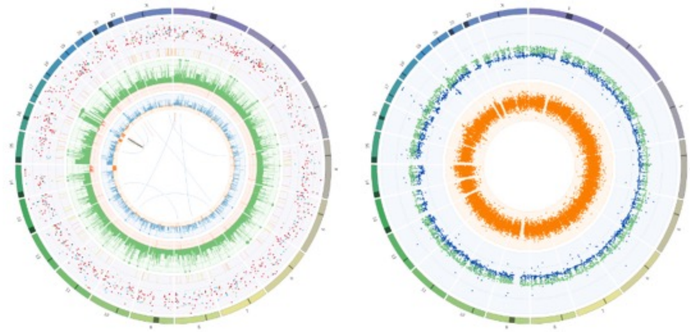

ORG70Org

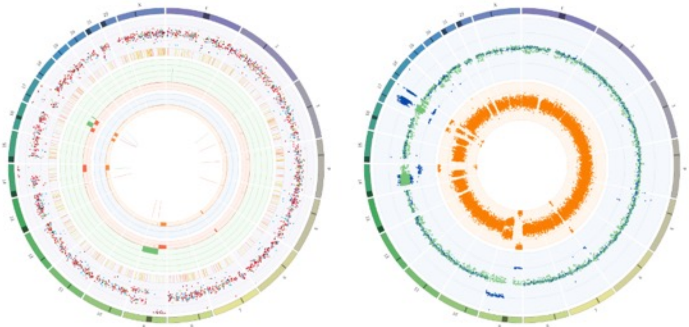

ORG73T

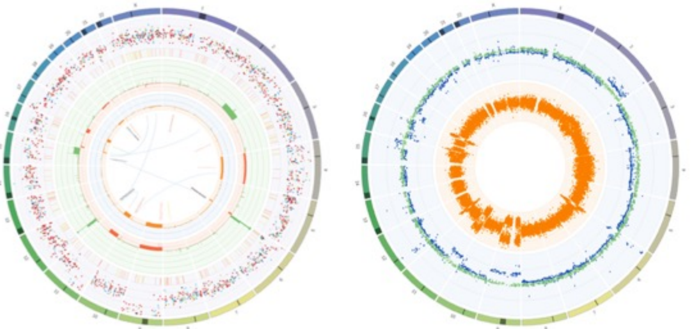

ORG73Org

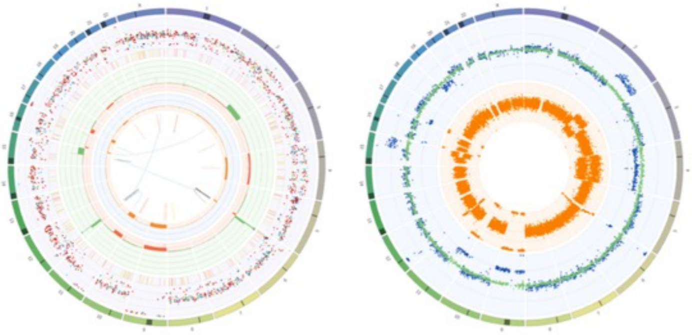

ORG74T

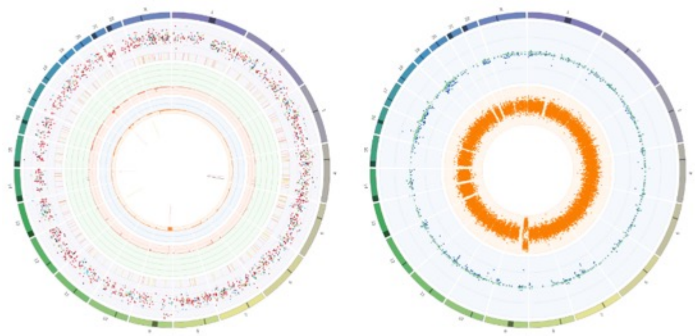

ORG74Org

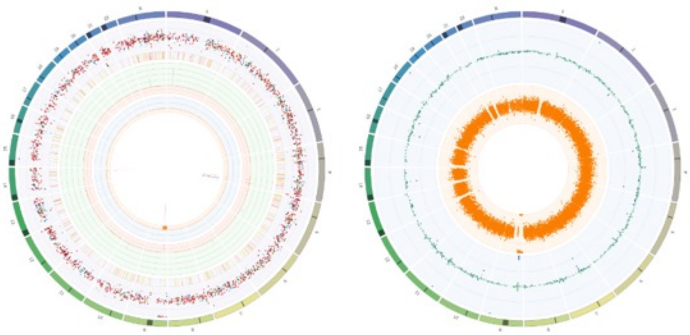

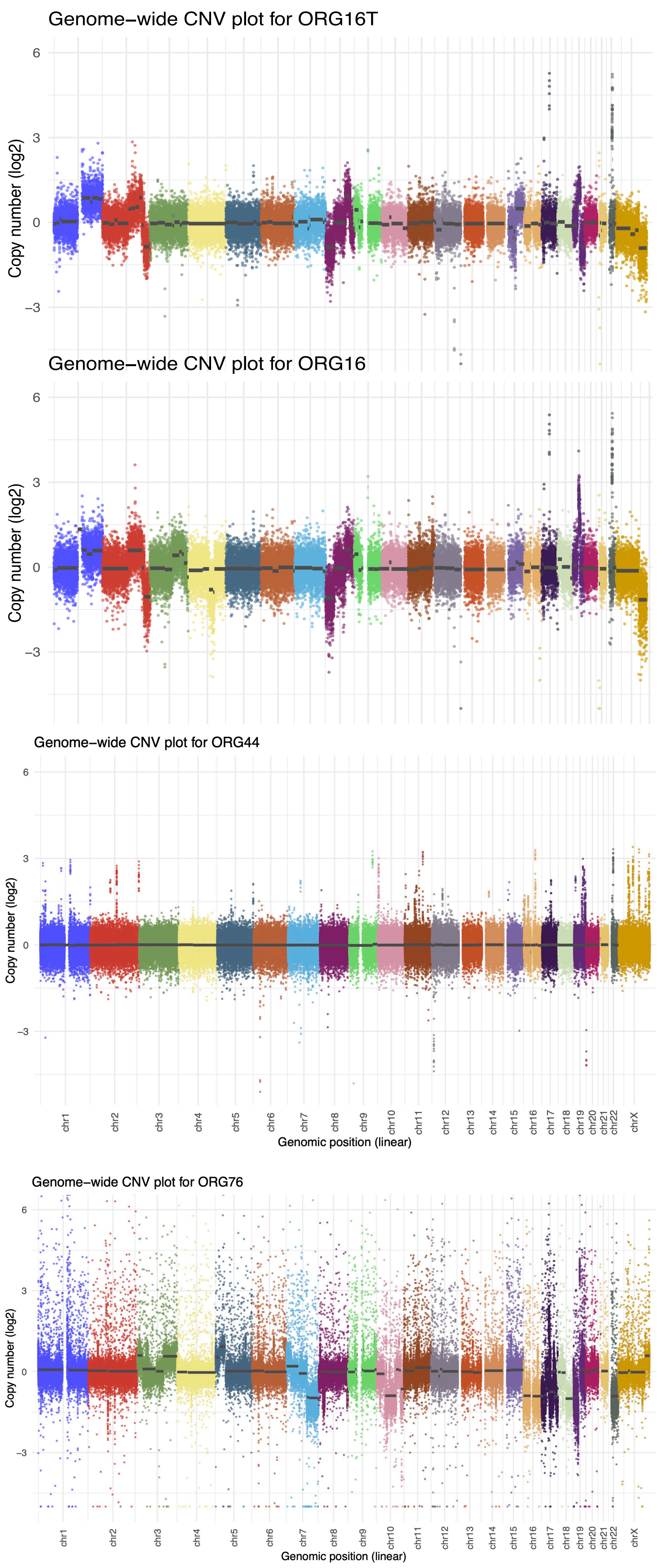


**Supplementary Figure 6 DNA sequencing circos and copy number plots**. Outputs from Oncoanalyser showing two circos plots for each sample analysed by whole genome sequencing. At left is the ploidy corrected copy number with structural variants. Green – copy number gains, red – copy number losses, blue – allelic imbalance, orange – loss of heterozygosity, dots – variant allele frequency. At right is the uncorrected copy number (blue) and B-allele frequency (orange) plots. For ORG16, output is from CopywriteR, for ORG44 output from PureCN and ORG76 output from CNVkit are shown.

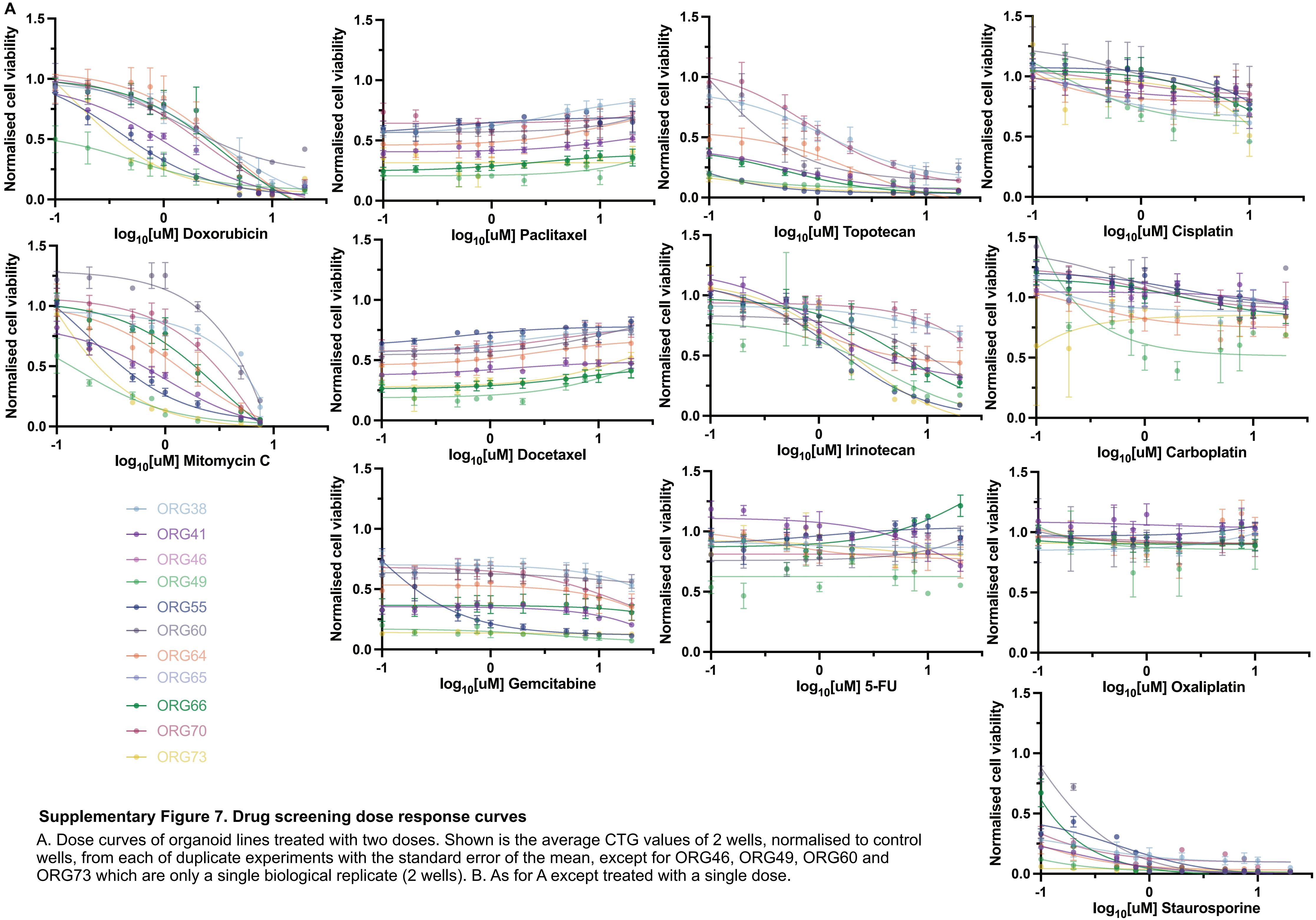


**Supplementary Figure 7. Drug screening dose response curves**

A. Dose curves of organoid lines treated with two doses. Shown is the average CTG values of 2 wells, normalised to control wells, from each of duplicate experiments with the standard error of the mean, except for ORG46, ORG49, ORG60 and ORG73 which are only a single biological replicate (2 wells). B. As for A except treated with a single dose.

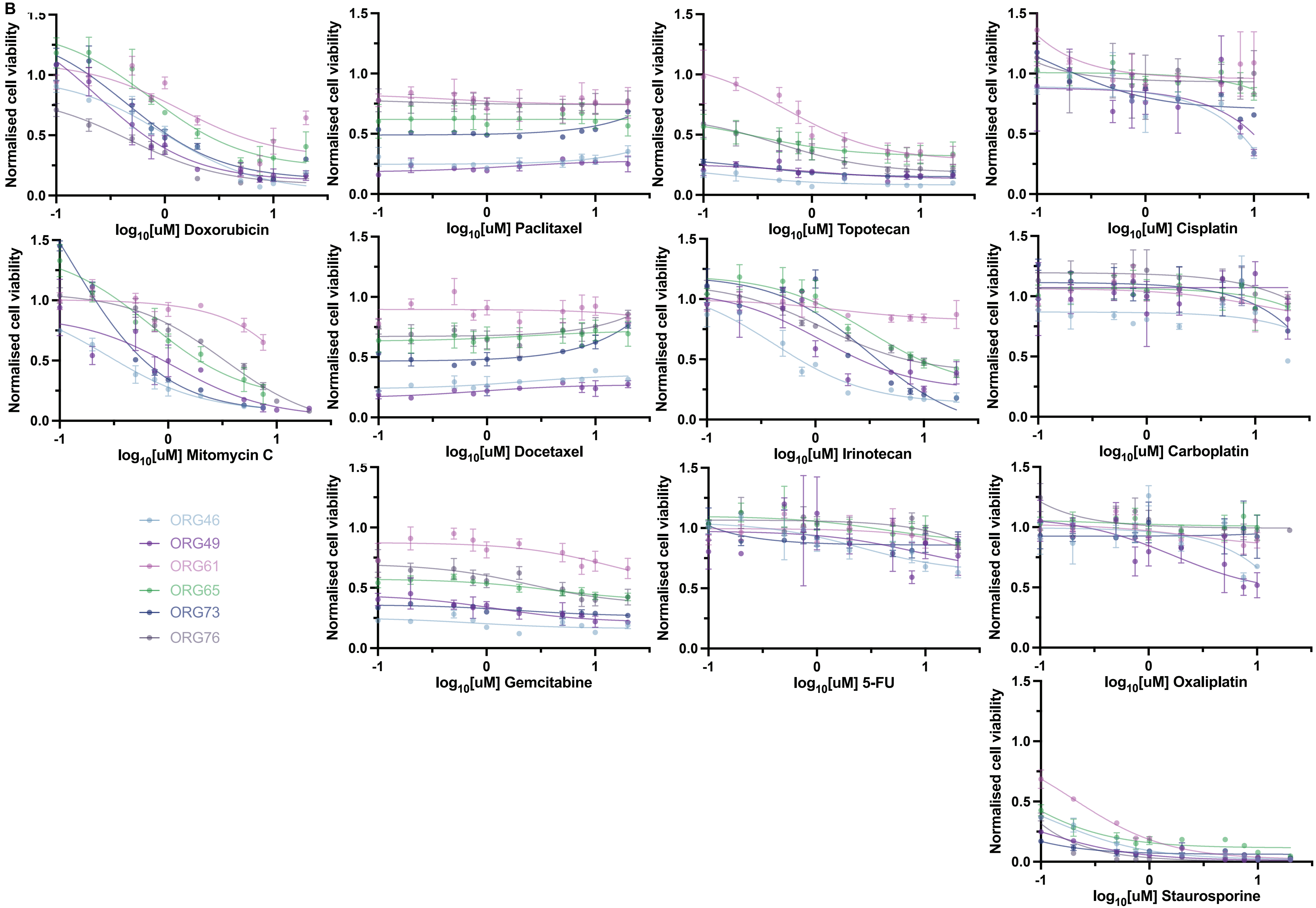

B
Normalised cell viability
log10[uM] Doxorubicin
log10[uM] Paclitaxel
log10[uM] Topotecan
log10[uM] Cisplatin
log10[uM] Mitomycin C
log10[uM] Docetaxel
log10[uM] Irinotecan
log10[uM] Carboplatin
log10[uM] Gemcitabine
log10[uM] 5-FU
log10[uM] Oxaliplatin
log10[uM] Staurosporine
ORG46
ORG49
ORG61
ORG65
ORG73
ORG76

### Supplementary Figure 8

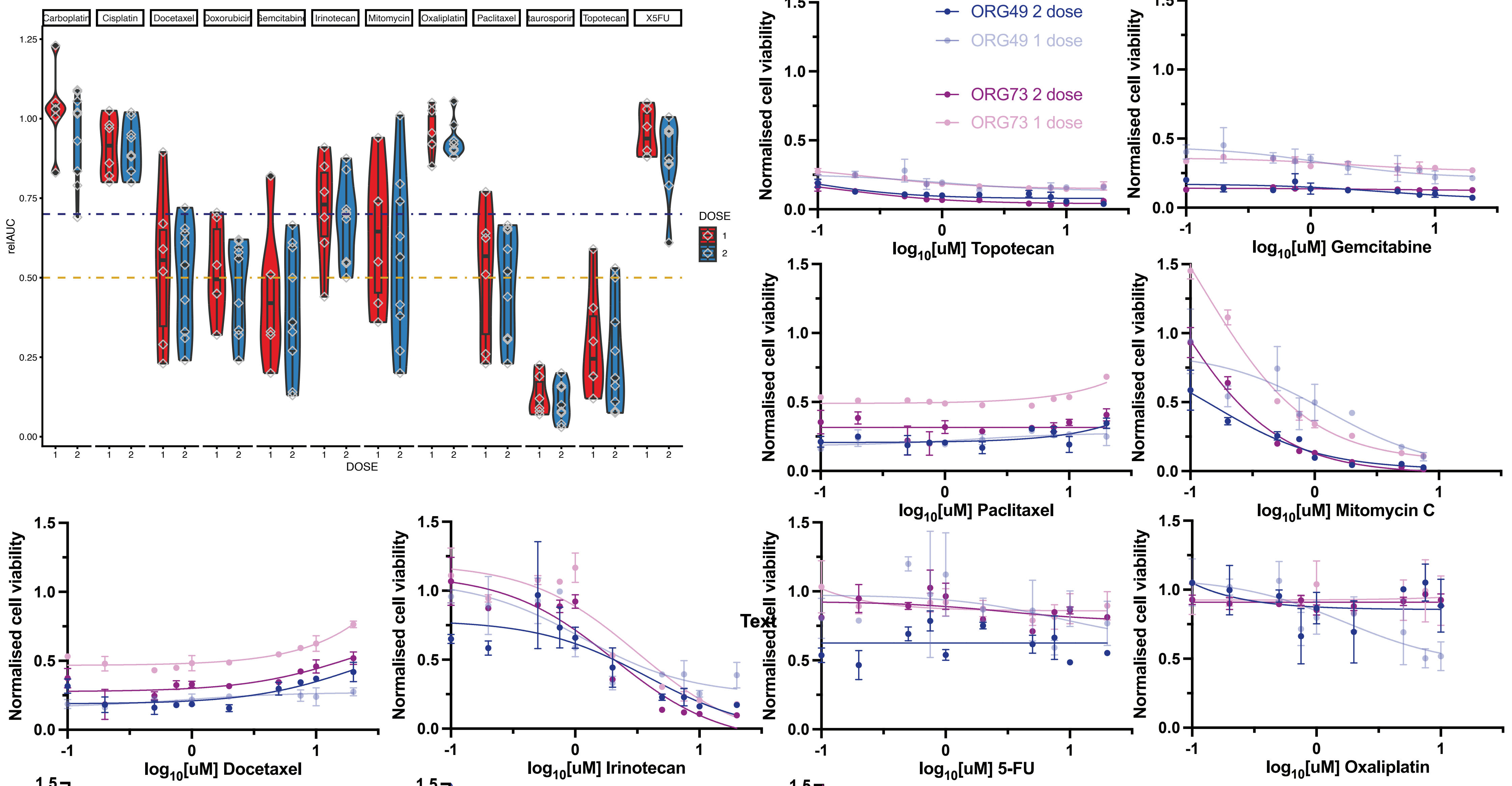


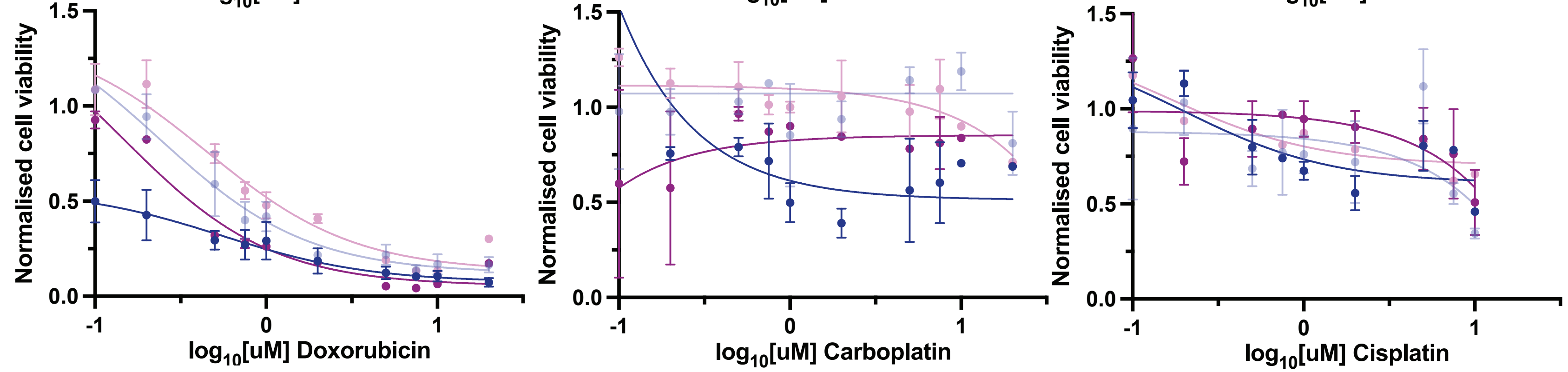


Supplementary Figure 8 Comparison between 1 and 2 dose experiments. Relative AUC of all cases, comparing 1 dose and 2 doses. Dose response curves for ORG49 and ORG73 for all chemotherapy drugs, which each had one experiment with a single dose and one with two doses. Error bars represent the range of two wells.

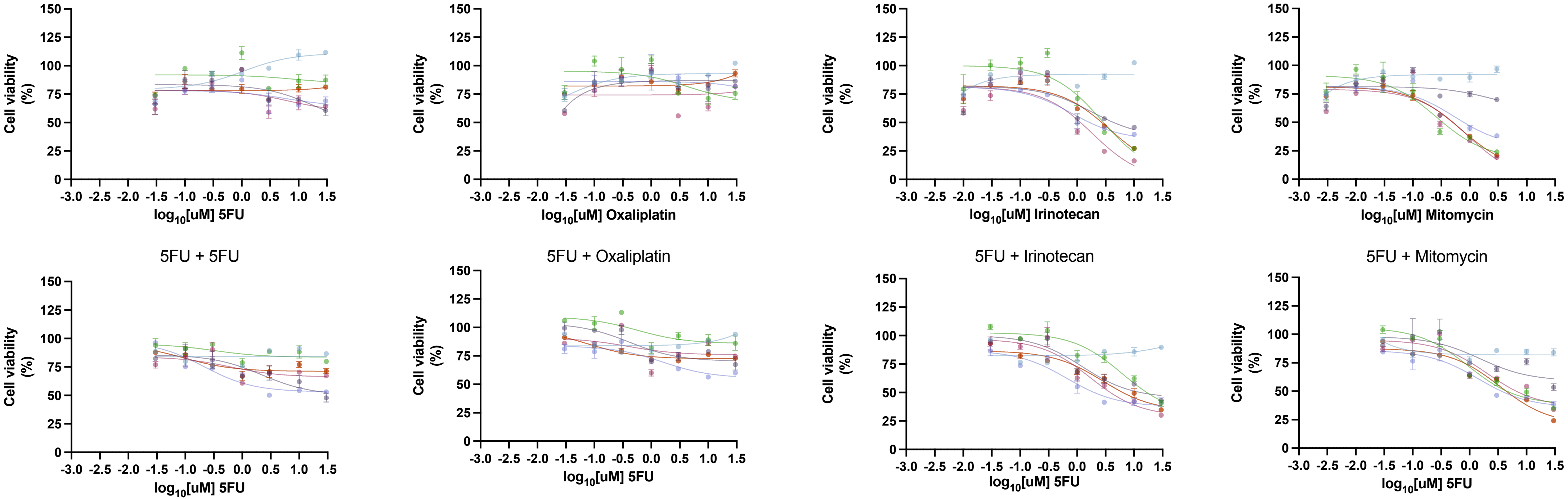


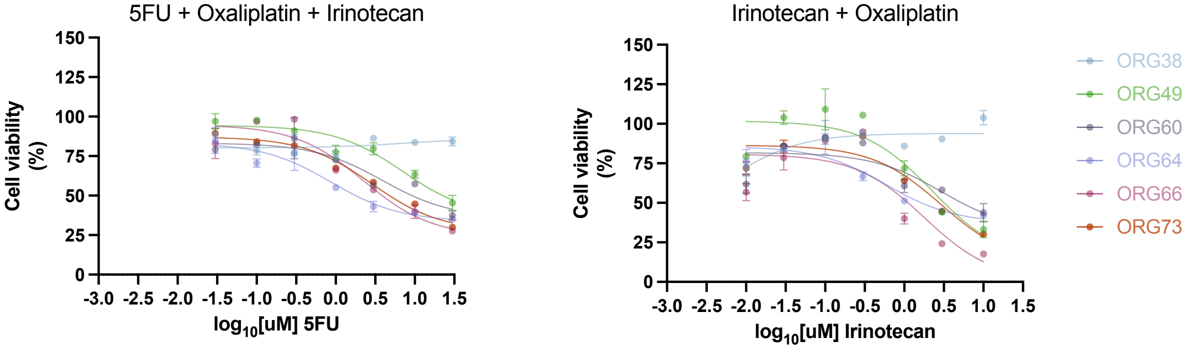


**Supplementary Figure 9. Synergy drug screens**

Cell viability as measured by CTG relative to control wells. Data points the average of two biological replicates, each the average of two wells. Error bars are the range

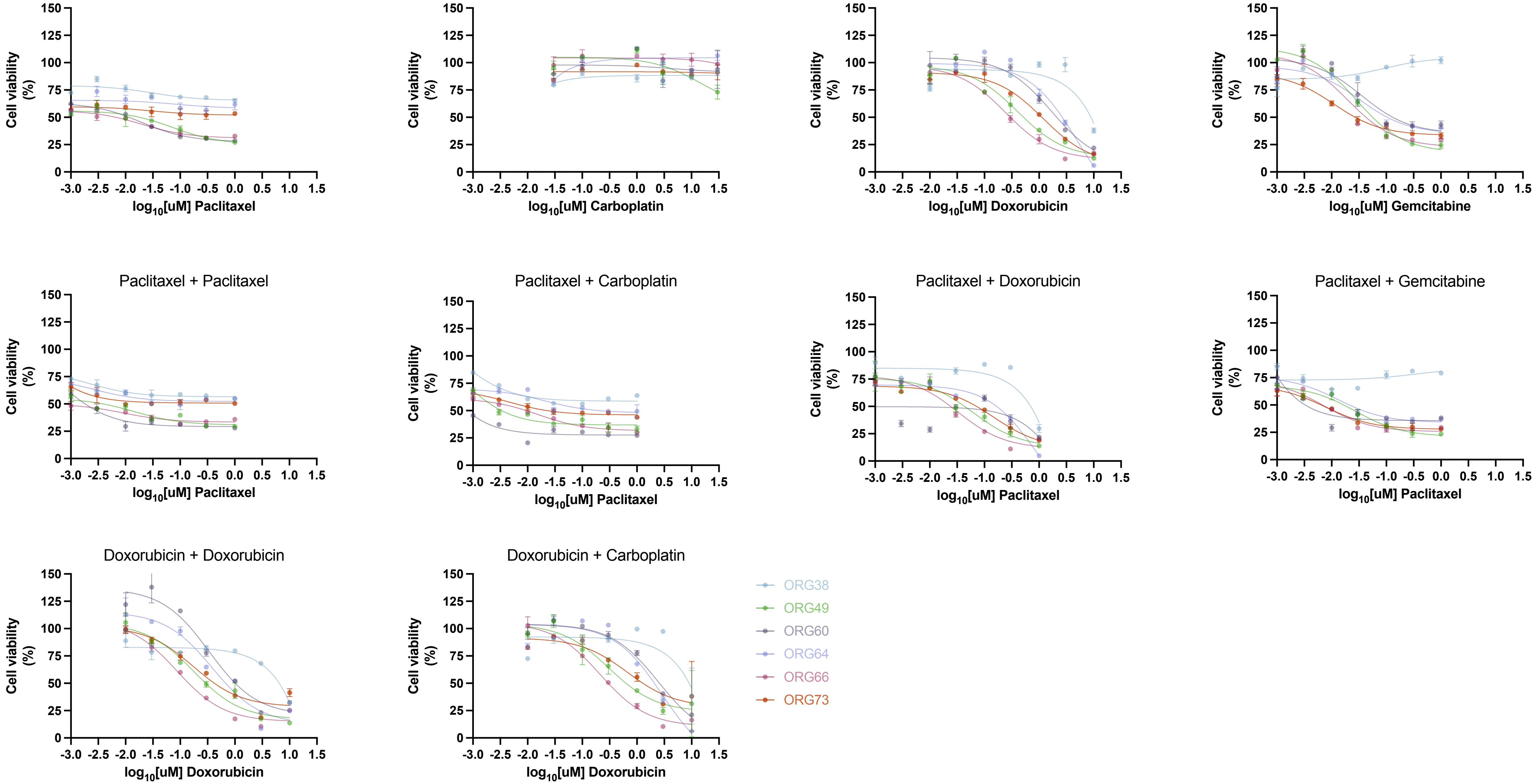

Cell viability (%)
log10[uM] Paclitaxel
log10[uM] Carboplatin
log10[uM] Doxorubicin
log10[uM] Gemcitabine
Paclitaxel + Paclitaxel
Paclitaxel + Carboplatin
Paclitaxel + Doxorubicin
Paclitaxel + Gemcitabine
Doxorubicin + Doxorubicin
Doxorubicin + Carboplatin
ORG38
ORG49
ORG60
ORG64
ORG66
ORG73

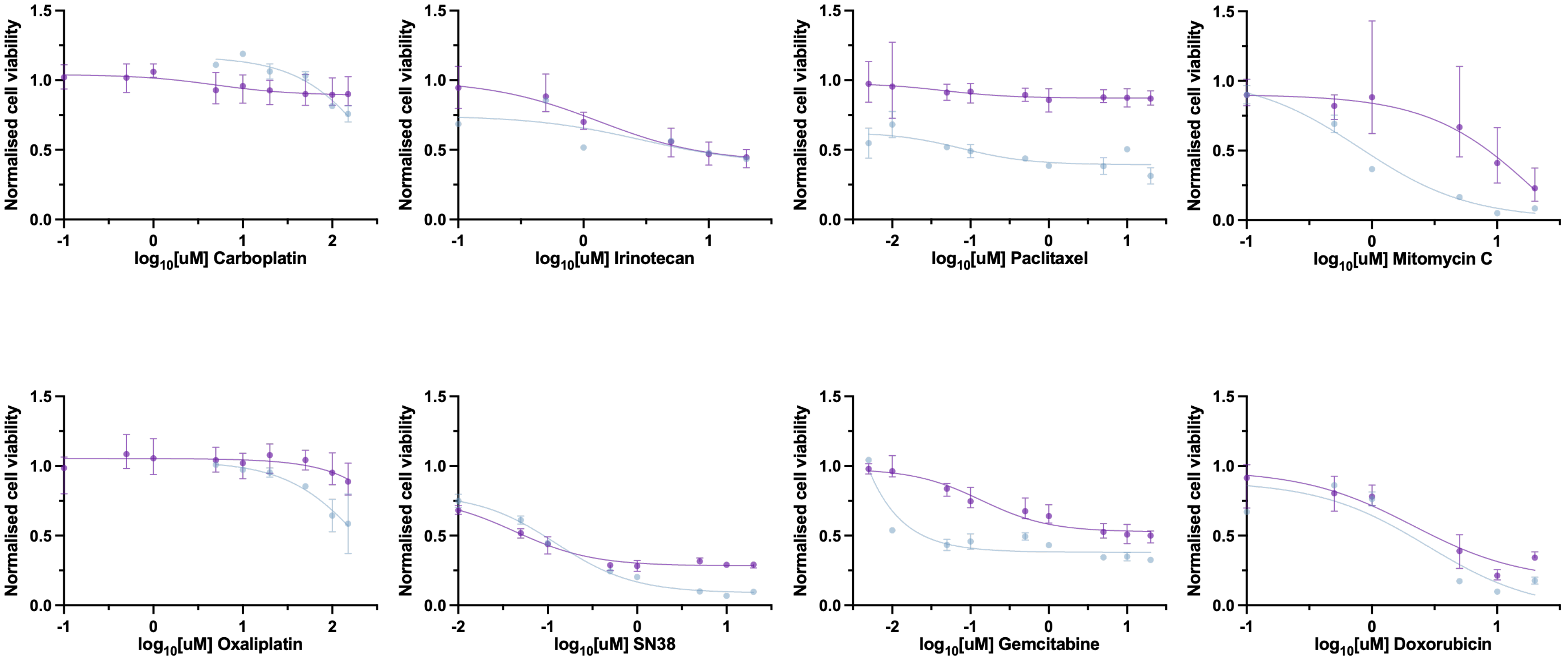


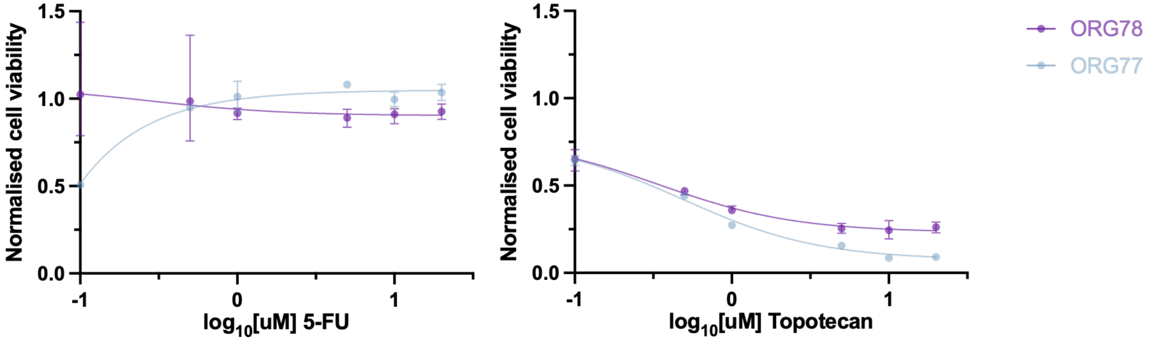


**Supplementary Figure 10. Early passage screening for clinical feasibility.** Dose response curves of CTG values normalised to control wells using a refined set of drugs compared to the previous screen (docetaxel and cisplatin removed, SN38 added). Note that additional lower doses were added for paclitaxel and gemcitabine, while higher doses were added for carboplatin and oxaliplatin. Average and range of two wells (ORG77) and four wells (ORG78 – two biological replicates)
