## Supplementary Table 2 for "Comprehensive drug efficacy data for mucinous ovarian carcinoma using a novel and extensive biobank of patient-derived organoid models"

|  | **CK7** | **p53** | **HER2** | **PAS** | **PAX8** | **CK20** | **Others** |
| --- | --- | --- | --- | --- | --- | --- | --- |
| **ORG16** | | | | | | | |
| Organoid | positive | abnormal positive | 3+ | positive | focal positive | negative | P16+ |
| Tissue section | positive | abnormal positive | 3+ | positive | focal positive | negative | P16+ |
| Path report | positive | abnormal positive | N/A | positive | focal positive | negative | p16+ CDX2 - WT1 patchy + |
| **ORG38** | | | | | | | |
| Organoid | positive | abnormal positive | negative (0) | positive | focal positive | negative | N/A |
| Tissue section | positive | abnormal positive | heterogeneous – invasive negative | positive | patchy positive | negative | N/A |
| Path report | positive | N/A | N/A | N/A | patchy positive | focal positive | ER- PR- CDX2 patchy + |
| **ORG41** | | | | | | | |
| Organoid | positive | WT | negative (0) | positive | negative | patchy positive | N/A |
| Tissue section | positive | WT | 1+ | positive | negative | focal positive | N/A |
| Path report | positive | N/A | N/A | N/A | positive | focal positive | CDX2 focal + |
| **ORG46** | | | | | | | |
| Organoid | positive | WT | negative (0) | positive | patchy positive | focal positive | N/A |
| Tissue section | positive | N/A | negative (0) | positive | patchy positive | focal positive | N/A |
| Path report | positive | N/A | N/A | N/A | N/A | patchy positive | SATB2: mostly –  P16 - |
| **ORG49** | | | | | | | |
| Organoid | positive | negative | 3+ | positive | negative | patchy positive | N/A |
| Tissue section | positive | abnormal negative | 3+ | positive | negative | negative | N/A |
| Path report | Glandular component +, solid - | abnormal negative | N/A | N/A | negative | Glandular component focal +, solid - | p16 patchy+ ER- PR- Glandular component CDX2+ CEA+; Solid component CDX2-CEA- Vimentin- AE 1/3 focal + synaptophysin focal + chromogranin- CD56- Ki67 variable, areas up to 80% + |
| **ORG55** | | | | | | | |
| Organoid | positive | abnormal positive | 1+ | positive | patchy positive | focal positive | N/A |
| Tissue section | patchy + | abnormal positive | 1+ | positive | patchy positive | focal positive | N/A |
| Path report | patchy + | N/A | N/A | N/A | negative | positive | MSH2+ MSH6+ MLH1+ PMS2+ ER - CDX2+ |
| **ORG59** | | | | | | | |
| Organoid | positive | weak positive >90% | 1+ | positive | focal positive | negative | N/A |
| Tissue section | positive | abnormal negative | 2+ | positive | patchy positive | patchy positive | N/A |
| Path report | positive | N/A | N/A | N/A | positive | positive | MSH2+ MSH6+ MLH1+ PMS2+ ER- PR- |
| **ORG60** | | | | | | | |
| Organoid | positive | abnormal positive | 1+ | positive | focal positive | negative | N/A |
| Tissue section | positive | abnormal positive | 2+ | positive | patchy positive | negative | N/A |
| Path report | positive | abnormal positive | 2+ | N/A | focal positive | negative | MUC5AC+ CDX2 focal + SATB2- MUC2- WT1-Napsin A- ER- PR- P16- |
| **ORG61** | | | | | | | |
| Organoid | positive | WT | negative (0) | positive | patchy positive | negative | N/A |
| Tissue section | positive | WT | 1+ | positive | negative | patchy positive | N/A |
| Path report | N/A | WT | N/A | N/A | N/A | N/A | N/A |
| **ORG63** | | | | | | | |
| Organoid | positive | abnormal positive | 1+ | positive | negative | negative | N/A |
| Tissue section | positive | WT | 1+ | positive | positive | negative | N/A |
| Path report | positive | WT | N/A | positive | focal positive | negative | CEA+ ER- PR- Vimentin- WT1- CDX2 focal + P16 patchy + MSH2+ MSH6+ MLH1+ PMS2+ |
| **ORG64** | | | | | | | |
| Organoid | positive | WT | negative (0) | positive | focal positive | negative | N/A |
| Tissue section | positive | WT | 1+ | positive | patchy positive | negative | N/A |
| Path report | positive | N/A | N/A | N/A | positive | focal positive | CEA+ ER - PR- Vimentin-  MSH2+ MSH6+ MLH1+ PMS2+ |
| **ORG65** | | | | | | | |
| Organoid | positive | abnormal positive | negative (0) | positive | negative | focal positive | N/A |
| Tissue section | positive | abnormal positive | 1+ | positive | focal positive | patchy positive | N/A |
| Path report | positive | WT | negative (0) | N/A | focal positive | focal positive | CDX2 patchy + SATB2- WT1- Napsin A- PR-  MSH2+ MSH6+ MLH1+ PMS2+ Ki67 within expected range |
| **ORG66** | | | | | | | |
| Organoid | positive | Weak nuclear positive >50% | 3+ | positive | focal positive | focal positive | N/A |
| Tissue section | positive | Weak nuclear positive >50% | 3+ | positive | focal positive | patchy positive | N/A |
| Path report | positive | N/A | N/A | N/A | focal positive | focal positive | CDX2 focal + ER- PR- WT1- MSH2+ MSH6+ MLH1+ PMS2+ |
| **ORG70** | | | | | | | |
| Organoid | positive | abnormal positive | 1+ | positive | negative | negative | N/A |
| Tissue section | positive | abnormal positive | 2+ | positive | focal positive | patchy positive | N/A |
| Path report | positive | N/A | N/A | N/A | focal positive | patchy positive | N/A |
| **ORG71*** | | | | | | | |
| Organoid | positive | Heterogeneous | 1+ | positive | patchy positive | focal positive | N/A |
| Tissue section | patchy positive | abnormal positive | 1+ | positive | focal positive | positive | N/A |
| Path report | moderate staining | non-specific heterogenous staining | N/A | N/A | negative | positive | CDX2 moderate + SATB2 moderate + ER- PR- |
| **ORG73** | | | | | | | |
| Organoid | positive | abnormal negative | 1+ | positive | patchy positive | focal positive | N/A |
| Tissue section | positive | abnormal negative | 2+ | positive | patchy positive | focal positive | N/A |
| Path report | positive | N/A | N/A | positive | N/A | focal positive | N/A |
| **ORG74** | | | | | | | |
| Organoid | positive | WT | negative (0) | positive | negative | patchy positive | N/A |
| Tissue section | positive | WT | 2+ | positive | focal positive | patchy positive | N/A |
| Path report | positive | N/A | N/A | N/A | negative | patchy positive | CDX2+ |
| **ORG76** | | | | | | | |
| Organoid | N/A | N/A | N/A | N/A | N/A | N/A | N/A |
| Tissue section | positive | N/A | negative (0) | positive | negative | negative | N/A |
| Path report | positive | N/A | N/A | N/A | negative | negative | GATA3 focal +. CDX2- TTF1- ER- PR- |
| **ORG77** | | | | | | | |
| Organoid |  |  |  |  |  |  |  |
| Tissue section |  |  |  |  |  |  |  |
| Path report | positive | WT | 2+ | N/A | positive | focal positive | CDX2 focal+ ER- PR- GATA3- CD10- Vimentin- TTF1- MSH2+ MSH6+ MLH1+ PMS2+ |
| **ORG78** | | | | | | | |
| Organoid |  |  |  |  |  |  |  |
| Tissue section |  |  |  |  |  |  |  |
| Path report | positive | WT | 2+ | N/A | patchy positive | patchy positive | CDX2 patchy+ ER- PR- MSH2+ MSH6+ MLH1+ PMS2+ |
